# Developmental constraints mediate the reversal of temperature effects on the autumn phenology of European beech after the summer solstice

**DOI:** 10.1101/2025.05.18.654771

**Authors:** Dominic Rebindaine, Thomas W. Crowther, Susanne S. Renner, Zhaofei Wu, Yibiao Zou, Lidong Mo, Haozhi Ma, Raymo Bucher, Constantin M. Zohner

## Abstract

Accurate projections of temperate tree growing seasons under climate change require representing developmental constraints that determine tree resource allocation. Recent work has identified a phenological “switch point” after the summer solstice (21 June), with pre-solstice warming advancing autumn phenology and post-solstice warming delaying it. Here, we propose this switch is flexible and occurs at the compensatory point between the antagonistic effects of early-season development and late-season temperature. We performed trans-solstice climate manipulation experiments on potted European beech (*Fagus sylvatica*) saplings to test (i) how spring leaf-out timing and June-August temperatures influence end-of-season timing (bud set and leaf senescence [50% loss of leaf chlorophyll content]), and (ii) whether daytime and nighttime temperatures before and after the solstice have different effects, given that trees primarily grow at night. Bud set and leaf senescence responses were tightly coupled (*R^2^* = 0.49), with bud responses being generally stronger. Each day delay in spring leaf-out delayed bud set by 0.24 ± 0.06 days and senescence by 0.22 ± 0.08 days. Post-solstice full-day cooling in July delayed autumn phenology in late-leafing individuals (bud set +4.9 ± 2.6 days; senescence +3.1 ± 2.8 days) but had negligible impact on early-leafing trees (bud set +1.4 ± 2.6 days; leaf senescence +2.2 ± 2.8 days). Conversely, August full-day cooling advanced both stages. Daytime cooling before the solstice had no effect, while after the solstice it advanced autumn phenology. Nighttime cooling always delayed bud set. These findings support the Solstice-as-Phenology-Switch model and highlight the central role of developmental progression in constraining growing seasons. Faster early-season development –especially under nighttime warming– moves trees past the switch earlier, increasing sensitivity to late-season cooling and thereby triggering earlier autumn phenology. To improve growing season length projections, phenology models should account for these developmentally-mediated and diel-specific temperature responses.

## Introduction

Climate shifts are leading to rapid, species-specific changes in phenology and ecosystem productivity (Boisvenue & Running, 2006; Menzel *et al*., 2006; Yang & Rudolf, 2010; Thackeray *et al*., 2016). In temperate forests, changes in the timing of spring leaf-out, autumn leaf-senescence and bud set are modifying water, energy and carbon cycles (Peñuelas *et al*., 2009; Richardson *et al*., 2013), with extended growing seasons increasing net ecosystem carbon uptake by up to 9.8 gC m^-2^ day^-1^ (Keenan *et al*., 2014). Therefore, understanding the interaction between climate change and temperate forest phenology is pivotal to improving forecasts of community dynamics and carbon sequestration.

The past few decades have seen delays in the onset of temperate autumn phenology, but these changes are much smaller in magnitude compared to the advances in spring leaf-out observed during the same period (Gill *et al*., 2015; Piao *et al*., 2019). This is unexpected, given that experiments have demonstrated a high sensitivity of leaf senescence to autumn warming, with phenological responses even surpassing the temperature sensitivity (days per °C) of spring leaf-out (Fu *et al*., 2018). One possible explanation for this discrepancy is that other factors may counterbalance the effects of autumn warming, with some studies finding that earlier leaf-out leads to earlier leaf senescence (Fu *et al*., 2014; Keenan & Richardson, 2015; Zani *et al*., 2020). This connection between spring and autumn phenophases could be due to developmental and nutrient constraints that affect carbon source-sink dynamics in temperate trees (Paul & Foyer, 2001; Zani *et al*., 2020; Zohner *et al*., 2023; Gessler & Zweifel, 2024), imposing limits on the phenological growing season.

The Solstice-as-Phenology-Switch hypothesis, supported by experiments and observational data, posits that autumn phenology is driven by two counteracting temperature effects (Zohner *et al.,* 2023). From the start of the growing season, warmer air temperatures drive faster development and growth activity, such as tissue hardening, meristematic activation, cell division and maturation, allowing trees to fulfil their developmental requirements more quickly and initiate leaf senescence earlier (Körner, 2021; Tumajer *et al*., 2021; Körner *et al*., 2023). The majority of activity occurs well before the end of the phenological growing season (Cruz-García *et al*., 2019; Etzold *et al*., 2022; Körner *et al*., 2023), so this effect is expected to act primarily during the early-season (henceforth referred to as the early-season developmental [ESD] effect). Conversely, the rate at which senescence progresses is predominantly mediated by temperature (Estiarte & Peñuelas, 2014; Fu *et al*., 2018; Zohner *et al*., 2023), with cooling triggering senescence progression and thus, warmer temperatures slowing senescence and delaying the end of the growing season (henceforth referred to as the late-season temperature [LST] effect). Year-round warming will therefore exert non-linear effects, with early-season cooling delaying autumn phenology (ESD effect), while late-season cooling advances it (LST effect). This seasonal reversal in temperature responses provides a mechanistic explanation for why shifts in autumn phenology are typically smaller and less consistent in direction than those observed for spring leaf-out (Piao *et al*., 2019). The fact that this reversal occurs after the solstice points to the role of photoperiod in regulating plant physiology (Bauerle *et al*., 2012; Petterle *et al*., 2013; Singh *et al*., 2017). However, the realised timing of this reversal is context-specific and appears to have advanced in recent decades (Zohner *et al*., 2023).

To explain this flexibility, we propose that the solstice acts as an environmental switch, with declining daylength providing a consistent and biologically meaningful cue that initiates the LST effect, while the ESD effect can persist beyond the solstice depending on an individual’s developmental state in a given year. Under this framework, the reversal in temperature responses occurs at a compensatory point where the advancing ESD effect is balanced by the delaying LST effect (Fig. 1). This point should therefore not be fixed to a calendar date but instead vary with developmental progression each year.

**Figure 1.**
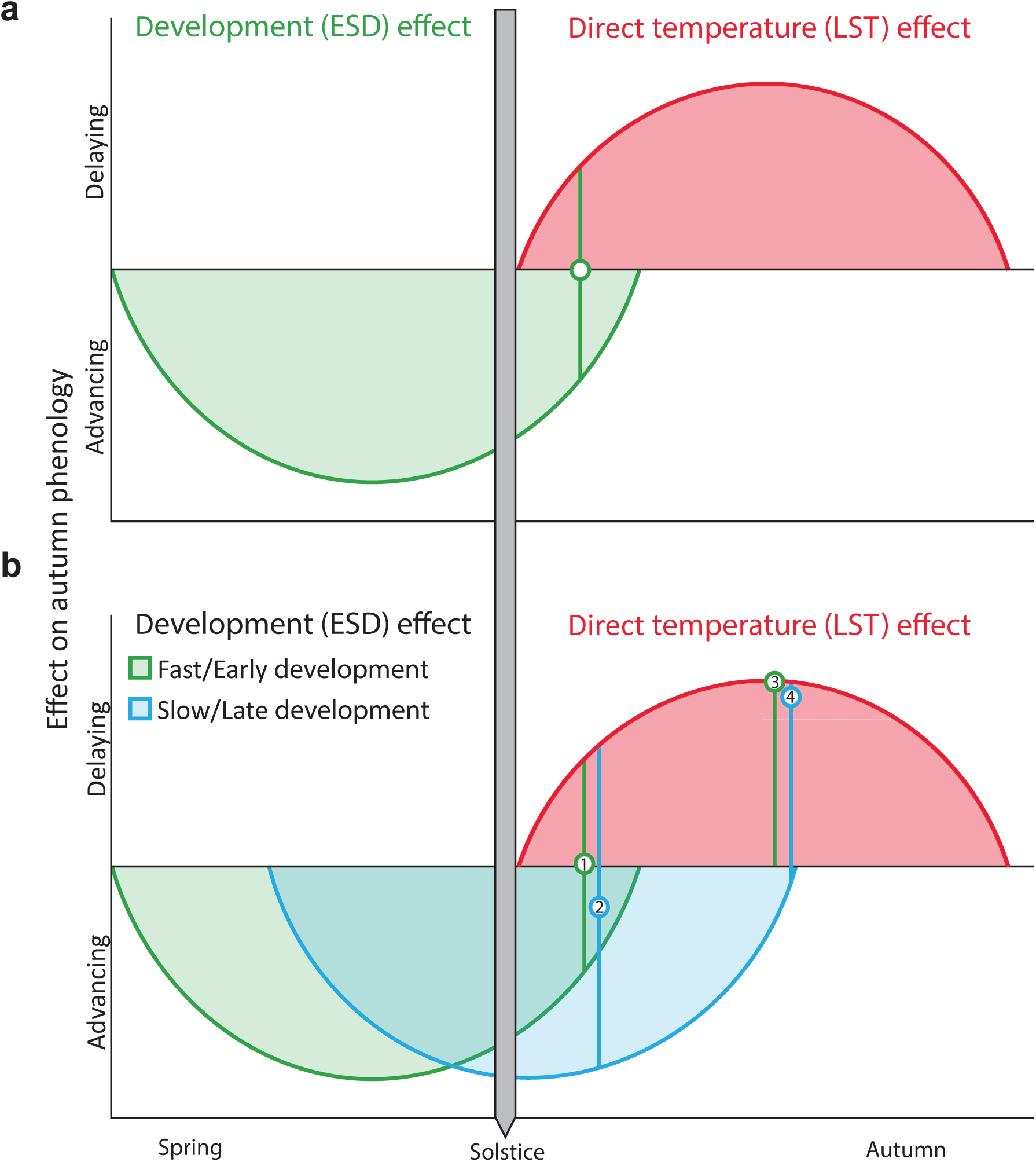
Conceptual model of autumn phenological responses of temperate trees to early-season development (ESD) and late-season temperature (LST) effects. Autumn phenology, represented in this study by the timing of primary growth cessation (bud set) and leaf senescence (50% loss of leaf chlorophyll content), is influenced by two opposing factors: ESD and LST. a) In our model, ESD, which is driven by temperature, has an advancing effect on autumn phenology that lasts until shortly after the summer solstice (green curve). Higher temperatures cause trees to complete their annual life cycles faster, allowing them to set buds and senesce leaves. After the summer solstice, as days shorten, trees become increasingly sensitive to cooling conditions, so late-season warming slows the progression of bud set and senescence, delaying autumn phenology (LST effect, red curve). The compensatory point, where the advancing ESD effect is balanced by the delaying LST effect, is represented by the green circle. b) According to the model, the position of this compensatory point is flexible and varies between years based on the speed of development. When development is slow or starts late (blue curve), the compensatory point is reached later than under fast or early development (green curve). Therefore, shortly after the solstice, the effect of temperature on autumn phenology differs between fast/early and slow/late developing individuals. For example, point 1 (green circle) shows no net temperature effect on phenology in fast/early trees. By contrast, point 2 (blue circle) shows that in slow/late developing trees, warmer temperatures shortly after the solstice (in July) still advance autumn phenology. However, as the growing season progresses and days shorten, trees become more responsive to cooling regardless of prior developmental speed (LST effect strengthens). Additionally, a weakening in the ESD effect is expected as individuals approach completion of their developmental requirements. By August, both fast/early and slow/late trees should therefore exhibit similar phenological responses, with warming consistently delaying autumn phenology (points 3 and 4).

For temperate trees, developmental outcomes are the production of viable seeds, cessation of primary and secondary growth, and maturation of perennating tissues including leaf and flower buds (i.e., completion of bud set) before the onset of frost (Rohde & Bhalerao, 2007; Tanino *et al*., 2010; Cooke *et al*., 2012). Because environmental conditions constrain the rate and duration of development, the timing of this compensatory point should vary among years and individuals. Under more advanced early-season development, this point is expected to be reached shortly after the solstice, whereas under slower development it occurs later (Fig. 1). However, such flexibility has yet to be demonstrated for processes directly linked to growth cessation, such as autumn bud set.

In this study, we investigate how air temperature changes around the summer solstice affect end-of-season timing in European beech (*Fagus sylvatica* L.), using bud set and leaf senescence as key physiological markers of autumn phenology (Singh *et al*., 2017; Zohner & Renner, 2019; Mariën *et al*., 2021). To elucidate the underlying mechanisms, we employ strong physiological forcing experiments specifically designed to maximise signal-to-noise ratios rather than representing contemporary or future climates. We ask three main questions: i) How does cooling before vs. after the solstice affect end-of-season timing? ii) How do diel (daytime vs. nighttime) temperature changes differ in their effects? iii) Does the timing of leaf-out modulate the reversal of temperature effects on autumn phenology? Diel temperature variations are relevant because trees typically grow more at night when temperatures and water deficit are lower (Steppe *et al*., 2015; Mencuccini *et al*., 2017; Zweifel *et al*., 2021). Thus, developmental effects mediated by temperature may be especially pronounced during nights. Moreover, long-term trends in climate show asymmetries between daytime and nighttime warming (Vose *et al*., 2005; Zhong *et al*., 2023). To address the third question, we experimentally manipulated spring conditions to create early-leafing and late-leafing individuals within the same growing season, allowing us to assess whether slowed early-season development postpones the point at which late-season cooling advances autumn phenology (Fig. 1). As trees continuously adapt their physiological responses to environmental conditions over time, we expected their temperature responses to differ both across months (June-August) and between day and night.

## Materials and Methods

### Selected species

*Fagus sylvatica* L. has a large temperate European distribution, is highly economically and ecologically important, and is likely threatened by climate change (Gessler *et al*., 2006). Therefore, improving our understanding of how *F. sylvatica*’s growing season is modulated is vital to the management of European forests. Moreover, *F. sylvatica* has frequently been used as a key species to study temperate tree phenology (Dittmar & Elling, 2006; Vitasse *et al*., 2011; Fu *et al*., 2018; Zohner & Renner, 2019; Zani *et al*., 2020; Zohner *et al*., 2023; Mariën *et al*., 2025; Švik *et al*., 2025).

### Experiment 1

To test the antagonistic influences of the ESDE and the LSTE on autumn phenology, we set up an experimental population (n = 267) of 40-60 cm tall European beech (*F. sylvatica*) trees in Zurich, Switzerland in 2023. The trees were sourced from a local nursery, and each tree was placed individually in a 20 L plastic pot containing a 1:1:1 sand / peat / organic soil mixture with a Nitrogen (N) concentration of ∼65 g m^-3^, a Phosphate (P_2_O_5_) concentration of ∼140 g m^-3^, and a Potassium (K_2_O) concentration of ∼400 g m^-3^. Individual trees were assigned randomly to one of 10 treatment groups (26 ≤ n ≤ 27). Treatment groups were cooled at different times of the year, using different cooling levels (Fig. 2, Table 1). To arrest spring development and thereby generate early-leafing and late-leafing individuals, half of the experimental population was placed outside under ambient conditions, while the other half was cooled in climate chambers from 4 April to 24 May. The chambers were set to a low of 2°C at night and a high of 7°C during the day, following a simulated day-night cycle of temperature and light availability (13 h photoperiod at ∼4,300 lux). These cold conditions were not intended to mimic natural European conditions but to strongly slow development without causing damage. Temperatures below 10°C substantially limit development and growth by slowing cell division and mitotic activity, with rates approaching zero as temperatures near 0°C (Körner, 2021; Tumajer *et al*., 2021). From 24 May to 21 June, all trees were kept outside under ambient conditions in a randomised block design. All trees were monitored to observe their individual leaf-out dates, which was defined as the date when > 50% of their leaves had unfolded, corresponding to BBCH15 (Capdevielle-Vargas *et al*., 2015).

**Figure 2.**
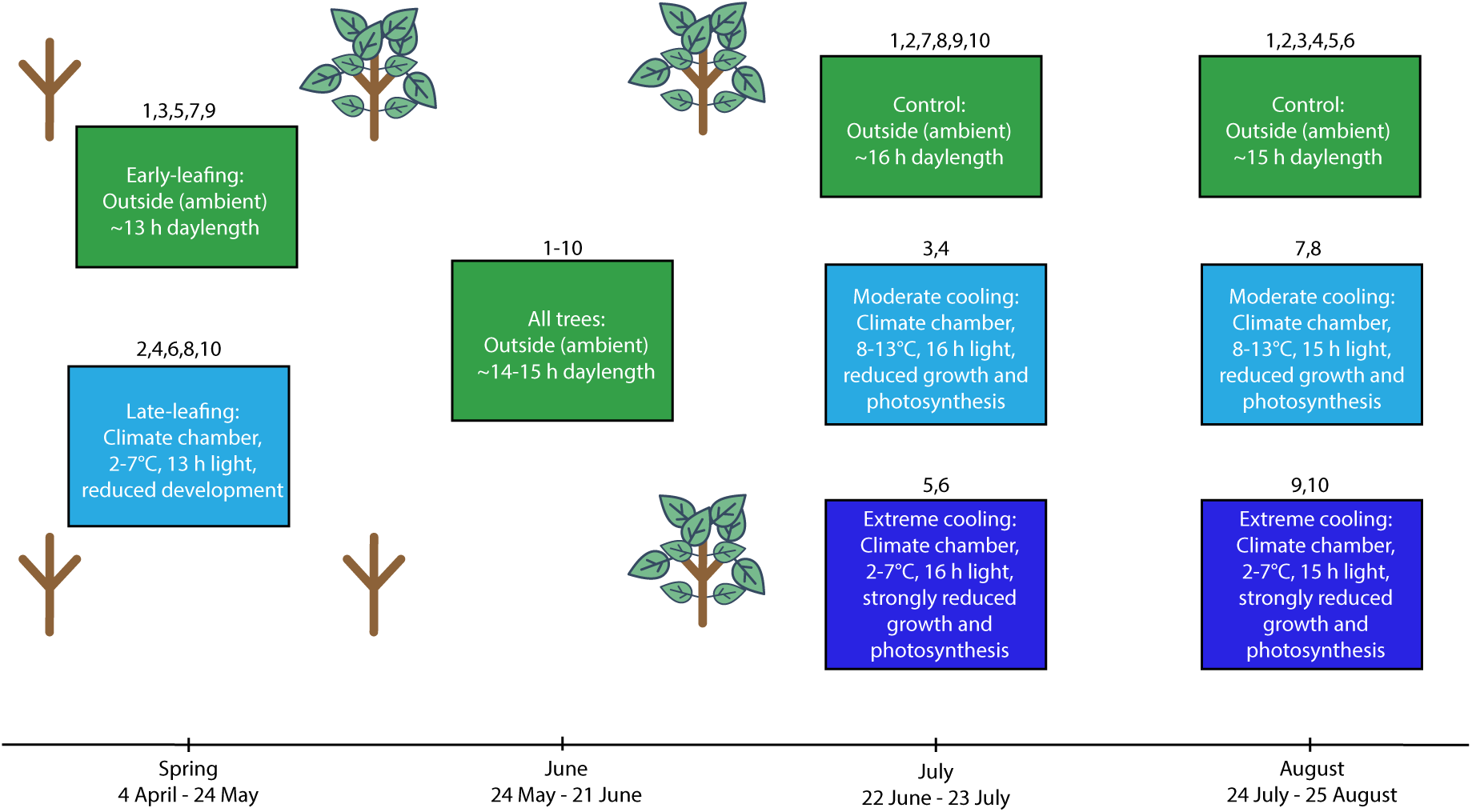
Depiction of the experimental timeline and settings for experiment 1. Each box corresponds to a specific treatment block at that point in time. The numbers inserted above the boxes refers to the treatment group (see Table 1 for details). Each box contains information on the location of the trees, the specific conditions they were under and the intended physiological effects of those conditions. The sapling graphics highlight differences in early-season developmental progression for the early-leafing and late-leafing groups. Following the August treatments all trees were placed outside under ambient conditions in a randomised block design.

**Table 1.** Description of the temperature treatments applied in experiment 1. July treatments were from 22 June to 23 July. August treatments were from 24 July to 25 August. Ambient means outside under natural conditions. Temperature ranges indicate the daily minimum and maximum temperatures experienced by trees inside climate chambers. The early-leafing and late-leafing trees were experimentally generated by placing potted trees in climate chambers from 4 April to 24 May and cooling them to 2°C at night and 7°C during the day to arrest spring development.

| Treatment | 1 | 2 | 3 | 4 | 5 | 6 | 7 | 8 | 9 | 10 |
| --- | --- | --- | --- | --- | --- | --- | --- | --- | --- | --- |
| Leaf-out timing | Early | Late | Early | Late | Early | Late | Early | Late | Early | Late |
| Treatment timing | - | - | July | July | July | July | August | August | August | August |
| Cooling level | Control (ambient) | Control (ambient) | Moderate (8-13°C) | Moderate (8-13°C) | Extreme (2-7°C) | Extreme (2-7°C) | Moderate (8-13°C) | Moderate (8-13°C) | Extreme (2-7°C) | Extreme (2-7°C) |

During the summer treatments, control trees were placed outside under natural ambient conditions. We did not use climate chambers for the control groups because warm conditions in these chambers can introduce unique physiological stressors—such as aphid infestations— not present in cold chambers (Bezemer *et al*., 1998). In experiment 2, however, we included an additional chamber control to test for potential chamber effects. Although we observed higher aphid abundances in these warm control chambers, chamber exposure itself had no detectable effect on autumn phenology. This strengthens the inference that the phenological shifts observed in our treatments reflect the intended temperature manipulations. Nevertheless, for experiment 1, where chamber controls were not included, we cannot entirely rule out the possibility that chamber conditions exerted some unintended influence on treated trees compared with ambient controls.

The July treatments took place directly after the summer solstice from 22 June to 23 July. Treated trees were placed in climate chambers set to a low of 2°C at night and a high of 7°C during the day or a low of 8°C at night and a high of 13°C during the day depending on their cooling level (extreme and moderate, respectively). Trees in the chambers experienced a photoperiod of 16 h at ∼7,300 lux. The extreme cooling was designed to severely impair cell division, expansion, and maturation as described above. Additionally, under this temperature regime, photosynthesis should be reduced by > 40% (Körner, 2006). The moderate cooling regime, which was still much colder than the ambient conditions, should have also impaired growth and development as well as reduced photosynthesis by at least 30% (Körner, 2006, 2021; Lenz *et al*., 2014). These settings aimed to generate large temperature differences between treatments and maximise our ability to detect the mechanism underlying the solstice switch, rather than to represent naturally occurring conditions. The August treatments took place from 24 July to 25 August under the same conditions as the July treatments except with a 15 h photoperiod. For the remainder of the experiment, all trees were kept outside in a randomised block design. Throughout the experiment, all trees were watered frequently to ensure constant water supply (Fig. S1).

To observe the effects of our treatments on the development of overwintering buds, we monitored bud growth to derive bud set dates, our marker for the cessation of primary aboveground development and the beginning of autumn phenology, for all trees. On each tree, the terminal bud of the primary shoot and the terminal bud on a random lateral stem were selected and tagged for measurement. Each selected bud was measured to 0.01 mm precision using a digital calliper (CD-P8”M, Mitutoyo Corp, Japan). We measured all buds on a regular basis from 4 July to 2 November. Bud set was defined as the date when each bud reached 90% of its own maximum length, which is considered to be an indicative stage of aboveground primary growth cessation (Signarbieux *et al*., 2017; Zohner & Renner, 2019). Bud lengths were linearly interpolated between measurement dates to derive the date at which they reached the 90% threshold.

To obtain a more holistic perspective of autumn phenology, we also derived leaf senescence dates from leaf spectral index measurements taken using a SPAD-502 Plus (Soil Plant Analysis Development, Minolta Camera Co., Ltd, Tokyo, Japan). We measured nine leaves per individual (three each from the top, middle and bottom of the crown) on a monthly basis in summer, every other week in September, and on a weekly basis from October to mid-December. We removed measurements taken on 7 December from the analysis as they were unreasonably high, likely due to false readings caused by extensive leaf browning. SPAD readings were then converted to total chlorophyll content (*Chl* in µg/g fresh weight) using an empirically established equation for *Fagus sylvatica* leaves (Percival *et al*., 2008):

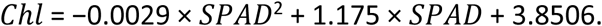

The chlorophyll content between two consecutive measurement dates was estimated using linear interpolation. For each treatment and measurement date, we removed individual chlorophyll measurements that were more than 1.5 × the interquartile range below the lower quartile or above the upper quartile. As an additional cleaning step, we completely removed any trees that had more than one data point removed in the last step. Finally, we calculated individual leaf senescence dates as the day-of-year when chlorophyll content last fell below 50% of the observed peak chlorophyll content.

### Experiment 2

To observe the effects of pre– and post-solstice daytime and nighttime temperature on autumn phenology, we set up an experimental population (n = 180) of four-year-old *Fagus sylvatica* trees in Zurich, Switzerland in 2022. The trees were sourced from a local nursery and each tree was placed individually in a 20 L plastic pot containing a 1:1:1 sand / peat / organic soil mixture with a Nitrogen (N) concentration of ∼65 g m^-3^, a Phosphate (P_2_O_5_) concentration of ∼140 g m^-3^, and a Potassium (K_2_O) concentration of ∼400 g m^-3^.

The ambient control treatment consisted of 36 trees exposed to natural ambient conditions. The remaining eight treatments were each applied to 18 trees and included cooling in climate chambers with simulated ambient day length (16 h) and light intensity (∼6,900 lux). The pre-solstice cooling treatments were applied between 22 May and 21 June. The post-solstice cooling treatments were applied between 22 June and 21 July. The pre– and post-solstice treatments included four levels each: Chamber control, where the trees were continuously subjected to 20°C; Day cooling, where trees were subjected to 8°C in the daytime and 20°C at night; Night cooling, where trees were subjected to 20°C in the day and 8°C at night; Full-day cooling, where trees were continuously subjected to 8°C (Fig. 3, Table 2). Due to warm temperatures, chamber control trees were subject to an increase in aphid abundance, however this did not alter their bud set timing (see data analyses). After treatment, all trees were placed in a randomised block design outside under ambient conditions. Soil moisture content was regulated by frequent watering. We measured all buds on a weekly basis from 25 August to 3 November following the same methodology as in experiment 1. We also measured leaf-level CO_2_ assimilation rates during the pre-solstice treatment window (see Zohner *et al.,* 2023 for methodology), and derived leaf senescence dates following experiment 1.

**Figure 3.**
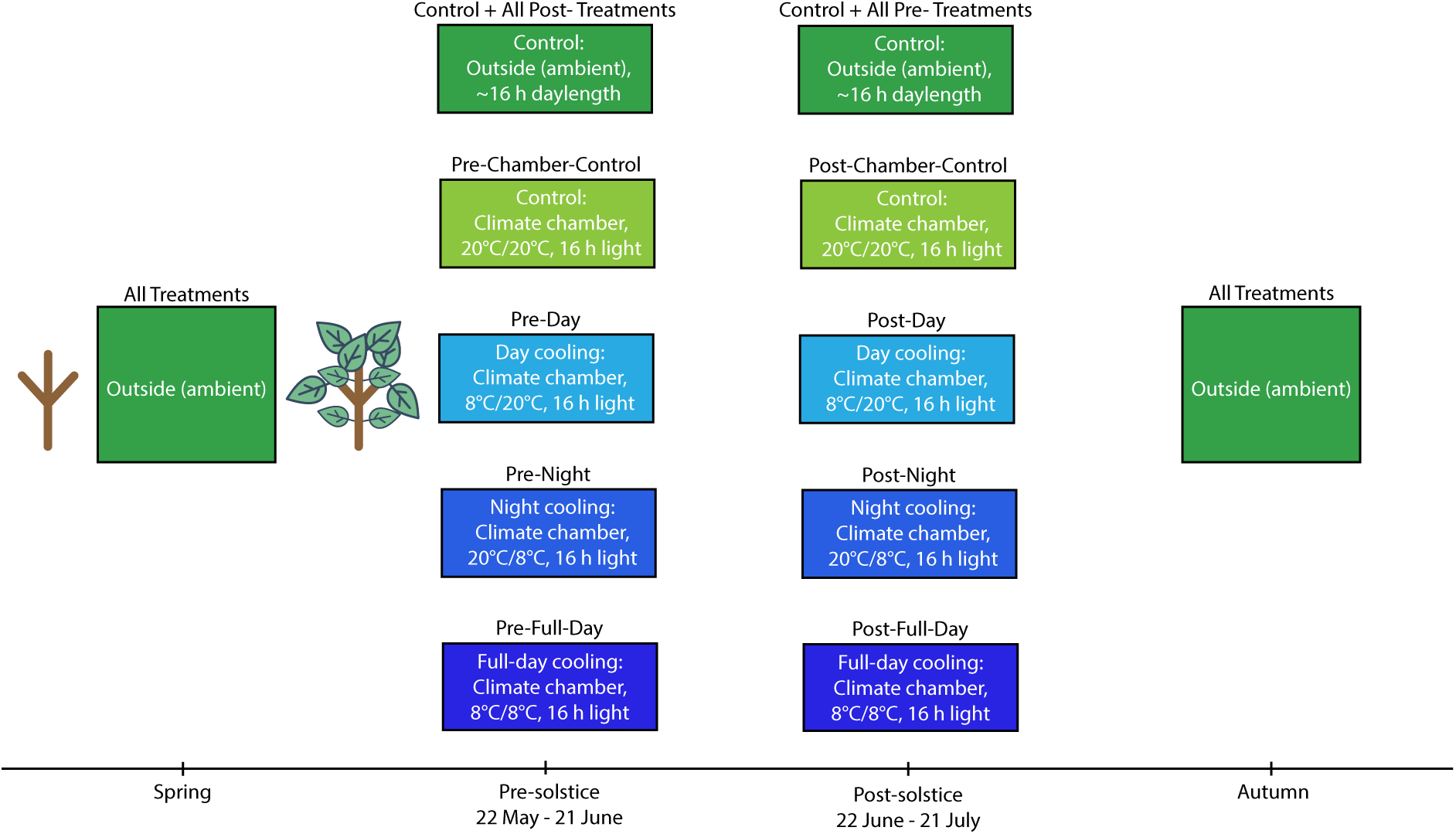
Depiction of the experimental timeline and settings for experiment 2. Each box corresponds to a specific treatment block at that point in time. The text inserted above the boxes refers to the treatment group (see Table 2 for details). Each box contains information on the location of the trees and the specific conditions they were under. The sapling graphics indicate that all trees had equal opportunities for early-season development.

**Table 2.** Description of the temperature treatments applied in experiment 2. June treatments were from 22 May to 21 June. July treatments were from 22 June to 21 July. Ambient means outside under natural conditions. The remaining temperature regimes are in the format day/night and refer to the temperatures applied to trees in climate chambers.

| Treatment | Control | Pre-Chamber control | Pre-Day | Pre-Night | Pre-Full-day | Post-Chamber control | Post-Day | Post-Night | Post-Full-day |
| --- | --- | --- | --- | --- | --- | --- | --- | --- | --- |
| Cooling time of year | - | June | June | June | June | July | July | July | July |
| Cooling time of day | - | - | Day | Night | Full-day | - | Day | Night | Full-day |
| Temperature regime | Ambient | 20/20°C | 8/20°C | 20/8°C | 8/8°C | 20/20°C | 8/20°C | 20/8°C | 8/8°C |

### Data analyses

For experiment 2, we performed one-way ANOVAs that showed no significant differences between the end-of-season dates of the ambient, pre-chamber and post-chamber control groups (for bud set: *F*_2,136_ = 0.346, p = 0.708, see Fig. S7; for leaf senescence: *F*_2,63_ = 0.931, p = 0.399). Therefore, for the following analyses we treated all three control treatments as one control treatment with 72 trees. For both experiments we ran linear models using treatment and bud type (apical vs. lateral) as predictors and bud set day-of-year, absolute bud growth or relative bud growth as the response variable. When modelling leaf senescence day-of-year, we used treatment as the sole predictor. For experiment 1, effect sizes were calculated in comparison to the corresponding control treatment, i.e., all early-leafing treatments were compared to the early-leafing control treatment, and all late-leafing treatments were compared to the late-leafing control. Absolute and relative bud growth were calculated as:

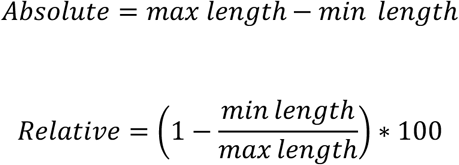

Where:

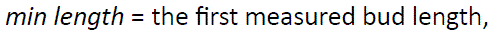

*max length* = the measured bud length when the bud first surpassed 90% of its final length. To determine the sensitivity of autumn bud set timing to spring leaf-out timing, we ran a linear mixed-effects model using leaf-out day-of-year and bud type as the fixed predictor variables, summer temperature treatment as random effect, and bud set day-of-year as the response variable. We ran the same analysis for leaf senescence timing without the bud type effect.

To quantify the relative contributions of the ESDE and the LSTE to variation in bud set timing in experiment 1, we ran a variance partitioning analysis. We fit a linear model with leaf-out day-of-year, summer cooling treatment (combination of cooling timing and level, e.g. July_Moderate), and bud type as explanatory variables. Variance partitioning was conducted using the *varpart* function in the *vegan* R package (Oksanen *et al*., 2025), which decomposes explained variance into unique and shared components. Additionally, we calculated the intra-class correlation coefficient (ICC) to compare inter-individual (within treatment) vs between treatment variance (Nakagawa et al., 2026).

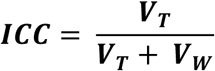

Where V_T_ = between treatment variance and V_W_ = within treatment variance. All statistical analyses were performed in R version 4.5.1 (R Core Team, 2025).

## Results

### Experiment 1

#### ESD effect

Linear modelling showed that across all treatments, late-leafing trees cooled during spring delayed their bud set by 4.6 ± 1.3 days (mean ± 2 SE, p < 0.01). Across all treatment pairings, the mean bud set date for the late-leafing group always occurred later than for the early-leafing group (Fig. S2). A linear mixed effects model including leaf-out day-of-year and bud-type as fixed effects and treatment as a random effect revealed that, on average, each day delay in spring leaf-out was associated with a delay of 0.24 ± 0.06 days in bud set timing. Across all treatments, bud type had a small effect, lateral buds set on average 1.1 ± 1.2-days earlier than apical buds (p = 0.06). When modelling only the early-leafing groups, bud type had no discernible effect (0.04 ± 1.64 days earlier for lateral buds, p = 0.96). However, when modelling only the late-leafing groups, lateral buds set earlier than apical buds (2.17 ± 1.64 days advancement, p = 0.01).

#### LST effect

The effect of July cooling differed strongly between the early– and late-leafing trees: Within the early-leafing group, moderate cooling in July lead to a small, non-significant delay in bud set compared to the ambient July treatment (1.4 ± 2.6 days, p = 0.29). July cooling had a much greater impact on late-leafing trees, leading to a 4.8 ± 2.6-day delay in bud set compared to the ambient July treatment in the late-leafing group (p < 0.01).

In contrast to July cooling, the effects of August cooling did not differ between early– and late-leafing trees, advancing bud set in all treatment groups (Fig. 4b). Cooling in August lead to a 4.5 ± 2.6-day (p < 0.01) and a 4.4 ± 2.6-day (p < 0.01) advancement in bud set for early– and late-leafing trees, respectively. The extreme cooling treatments showed similar patterns, although slightly less clearly (Fig. S3).

**Figure 4.**
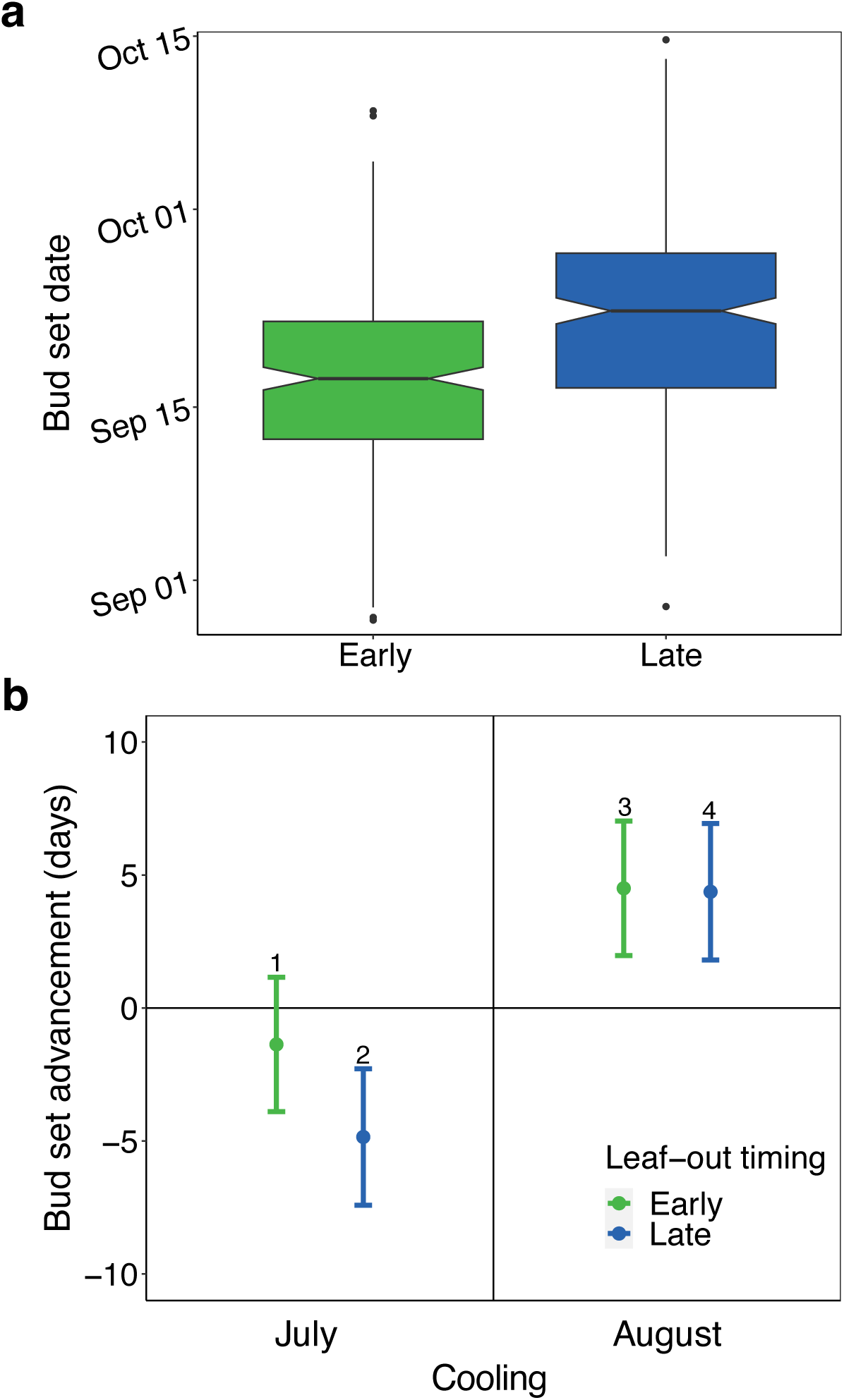
Effects of early-season development and late-season temperature on the timing of autumn bud set in *Fagus sylvatica* (experiment 1). **a**) Bud set dates for early– (green) and late– (blue) leafing trees including all treatments. Late-leafing trees were cooled (2-7°C) in climate chambers from 4 April to 24 May to arrest their development and delay leaf-out. **b)** Effects of July (22 June to 23 July) and August (24 July to 25 August) moderate cooling (8-13°C) on bud set date for early– (green) and late– (blue) leafing trees (see Fig. S3 for extreme cooling effects). Analyses show effect size means ± 95% confidence intervals from linear models, including treatment and bud-type (apical vs lateral) as predictors. Early-leafing effects are calculated against the early-leafing control and late-leafing effects are calculated against the late-leafing control. The bud type effect is not shown. Number labels (1-4) above each point are shown to aid comparison between points 1-4 in the conceptual model (Fig. 1b) and the observed effects. Positive values indicate advances in bud set and negative values indicate delays.

Variance partitioning revealed that the ESD effect (leaf-out day-of-year) explained 8.8% of the total variance in bud set timing, the LST effect (summer cooling treatment) explained 14.6%, and bud type explained 0.3%. Shared variance values were ≍ 0%, indicating no meaningful overlap. The intra-class correlation coefficient was 0.26, suggesting high levels of inter-individual (within treatment) variance. Effects of absolute and relative growth on bud set date can be found in the supporting information (Figs. S4-S5). Bud set timing had no effect on final bud length (Fig. S6).

### Experiment 2

Linear modelling showed that pre-solstice full-day (day and night) cooling delayed autumn bud set by 4.1 ± 3.6 days (mean ± 2SE, p = 0.02) (Fig. 5). Pre-solstice nighttime cooling had a similar effect, delaying bud set by 4.2 ± 3.6 days (p = 0.02). By contrast, pre-solstice daytime cooling had no significant effect on bud set (0.1 ± 3.7 days, p = 0.94). Post-solstice full-day cooling advanced autumn bud set by 5.2 ± 3.6 days (p < 0.01). Similarly, post-solstice daytime cooling advanced bud set by 5.3 ± 4.3 days (p = 0.01). Conversely, post-solstice nighttime cooling delayed bud set by 3.8 ± 3.5 days (p = 0.03). Across all treatments, lateral buds set considerably earlier than apical buds (9.4 ± 2.08 days, p < 0.01). Effects of relative growth on bud set date can be found in the supporting information (Figs. S8).

**Figure 5.**
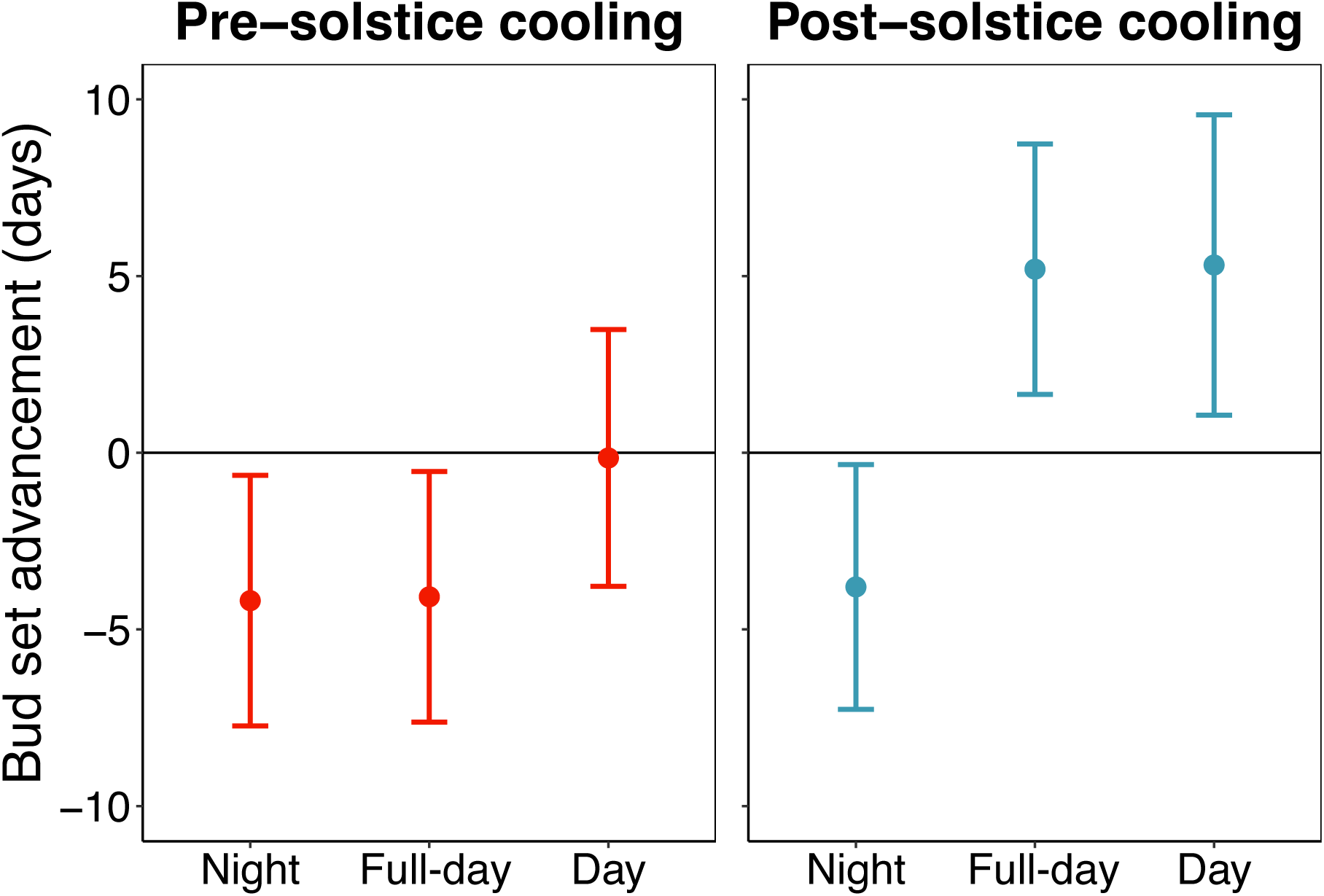
The pre– and post-solstice effects of night, full-day and day cooling on the timing of autumn primary growth cessation in *Fagus sylvatica* (experiment 2). Effects of pre-solstice (22 May to 21 June) and post-solstice (22 June to 21 July) cooling on bud set date. Full-day cooling trees were continuously cooled to 8°C, day cooling trees were cooled to 8°C in the day and kept at 20°C at night, night cooling trees were cooled to 8°C at night and kept at 20°C in the day. Analyses show effect size means ± 95% confidence intervals from linear models, including treatment and bud-type (apical vs lateral) as predictors. The bud type effect is not shown.

### Comparison of phenological metrics

Analyses using leaf senescence (50% drop in chlorophyll content) as the phenological marker produced results that were overall consistent with the bud set analyses. For example, in experiment 1, senescence was delayed by 4.22 ± 1.61 days in late-leafing trees and each day delay in spring leaf-out led to a 0.22 ± 0.08-day delay in leaf senescence, closely matching with bud set. Across both experiments the observed patterns were largely the same, although the strengths of each effect differed slightly, and overall, the effects on leaf senescence tended to be weaker (Figs. S9-10). Out of all treatments, opposing responses from bud set and leaf senescence timing were only observed in the early-leafing extreme August cooling group (Treatment 9, bud effect = 1.9 ± 2.59 days advancement, p = 0.14, leaf effect = 1.77 ± 3.53 days delay, p = 0.32) from experiment 1, and the post-full-day cooling group from experiment 2 (bud effect = 5.2 ± 1.81 days advancement, p < 0.01, leaf effect = 1.66 ± 3.34 days delay, p = 0.32). Out of 66 pairwise comparisons of estimated marginal means (every treatment compared against every other treatment in the same experiment), 55 (83%) showed directional agreement between bud set and leaf senescence effects (Fig. S11, adjusted *R^2^* = 0.49).

## Discussion

### Experimental forcing and scope of inference

The temperature manipulations applied in this study were intentionally designed as physiological forcing treatments rather than as simulations of realistic climatic scenarios. By imposing strong and temporally discrete constraints on development and sensitivity to cooling, we tested whether these physiological processes could be forced to advance or delay the timing of either autumn phenology, or the compensatory point, as predicted by our conceptual model (Fig. 1). Importantly, the exact date upon which the compensatory point was crossed was not (and with current knowledge cannot be) directly measured; instead, its position was inferred from changes in the direction and magnitude of temperature effects on autumn phenology before the solstice (June), directly after it (July), or later in the growing season (August). Accordingly, the compensatory point should be interpreted as an inferred conceptual node that structures seasonal temperature responses, rather than as a directly observed phenological event. Given the large individual variation expected in phenological experiments, we used single experimental populations of single provenance beech saplings to minimise uncontrolled for variation arising from genetic differences (Meger *et al*., 2021). The primary inference of these experiments concerns causality and constraint, i.e., whether these processes are capable of producing the observed patterns, rather than quantitative predictions of ecosystem responses to contemporary or future climates.

Our experiments on European beech test how bud set—a marker of primary growth cessation—and leaf senescence respond to monthly and diel temperature changes around the summer solstice. We found large differences in the responses between trees subjected to daytime versus nighttime cooling before and after the solstice (Fig. 5) and between early– and late-leafing trees (Fig. 4). These differences suggest that the ESD effect not only influences the timing of autumn phenology *per se* but also plays a critical role in determining the actualised timing of the reversal of phenological responses to temperature after the summer solstice. Therefore, our findings support our conceptual model (Fig. 1), highlighting development as a key factor underpinning the Solstice-as-Phenology-Switch hypothesis. In the following sections, we discuss the effects of pre-solstice and post-solstice air temperature on development, along with the distinct impacts of daytime versus nighttime temperature variations.

### Early-season development rates alter responses to late-season temperature

Our trans-solstice climate manipulation experiments showed that trees delayed the end of their growing season in response to slow/late early-season development (delayed spring leaf-out; ESD effect). Regardless of the summer cooling treatments, late-leafing trees consistently set buds and senesced leaves after their early-leafing counterparts (Fig. S2), with each day delay in spring leaf-out delaying bud set by an average of 0.24 days and senescence by 0.22 days. This is in line with previous studies demonstrating a tight linkage between within-year variations in spring and autumn phenology (Fu *et al*., 2014; Keenan & Richardson, 2015; Signarbieux *et al*., 2017), likely governed by developmental constraints (Zohner *et al*., 2023), buildup of water and nutrient stress (Paul & Foyer, 2001; Buermann *et al*., 2018; Bigler & Vitasse, 2021), or leaf aging (Lim *et al*., 2007).

Whilst lateral buds tended to set earlier than apical buds, the effect was not consistent across leafing groups, suggesting that bud position interacts with developmental context rather than exerting a uniform effect. This inconsistency may reflect differences in non-structural carbohydrate distribution and hormonal gradients among individuals at different developmental stages (Powell, 1988; Barbier *et al*., 2017; Singh *et al*., 2022). Under reduced early-season development, earlier lateral compared to apical bud set suggests enhanced apical dominance with increased allocation of resources to the sapling’s primary shoot.

Summer cooling treatment [LST effect] explained more variance in bud set timing than leaf-out day-of-year [ESD effect] (Adjusted *R^2^* = 0.15 and 0.09, respectively), with a clear interaction between the two effects. July cooling induced a delay in bud set dates 3.4 times greater in late-leafing trees compared to early-leafing ones (4.8 versus 1.4 days delay), which agrees with the expectations derived from our conceptual model (Fig. 1). This dependence of autumn phenological responses to summer cooling on developmental progress demonstrates flexibility in the compensatory point between the antagonistic influences of the ESD and LST effects. Although we did not directly identify a discrete timing at which this compensatory point occurred, the pronounced difference in July treatment effects between early– and late-leafing trees suggests a shift in the seasonal window during which cooling delays, rather than advances, autumn phenology. Together, these results support a development-dependent flexibility in the effective timing of the reversal in temperature responses, consistent with a flexible compensatory point governed by developmental progression.

In natural systems, these responses may arise from interannual variation in early-season development, leading to year-to-year differences in the timing at which cool temperatures switch from delaying autumn phenology to advancing it. This would offer a physiological explanation for the advancement in the response reversal observed between 1966 and 2015 in *Fagus sylvatica, Aesculus hippocastanum, Quercus robur, and Betula pendula* (Zohner *et al*., 2023). However, extrapolations to more complex natural ecosystems should be made with caution as our experimental design prioritised mechanistic inference over generalisability and predictive power. In line with this focus, summer cooling treatment (LST effect), leaf-out date (ESD effect) and bud type together explained a substantial, though not majority, proportion of the variation in bud set timing (24%). Several physiological mechanisms could, in principle, contribute to this unexplained variation. For example, declining photosynthetic assimilation has been linked to accelerated senescence (Krieger-Liszkay *et al*., 2019). However, experimental reductions of August photosynthetic rates in beech (52–72%), achieved through either cooling or shading, demonstrated that photosynthetic assimilation was not a driver of phenological responses in beech (Zohner *et al*., 2023). This suggests that photosynthesis is unlikely to explain the residual variation observed here.

Rather than missing explanatory variables, much of the remaining variation is attributable to individual-level differences, as suggested by the relatively low intra-class correlation coefficient [0.26] (Nakagawa *et al*., 2026). Individual variation likely reflects genetic diversity— even within a single provenance population—and epigenetic mechanisms that influence phenological timing (Crawley & Akhteruzzaman, 1988; Scotti *et al*., 2016; Carneros *et al*., 2017; Solvin & Steffenrem, 2019; Malyshev *et al*., 2022). Additional sources of within-treatment variability may include measurement error, microclimatic heterogeneity, and baseline differences among individuals although steps were taken to minimise these throughout the experiment. However, because control trees did not experience chamber time, we cannot entirely rule out some influence of this factor on our comparisons. In this context of high individual variability, leaf-out timing (ESD effect) and summer cooling treatment (LST effect) together explaining 23.4% of variation in bud set timing demonstrates the mechanistic importance of these processes.

August cooling induced comparable advances in bud set timing in both early– and late-leafing trees (4.4-4.5 days). The diminishing difference in cooling responses between the two groups from July to August suggests that, by August, phenology is primarily governed by the LST effect (see direct temperature effect in Fig. 1b). The observed increase in the LST effect from July to August may be driven by a reduction in the influence of the ESD effect or by declining daylength (Kramer, 1936; Petterle *et al*., 2013; Singh *et al*., 2021), but more likely by an interaction between the two. Firstly, a weakening in the ESD effect is expected as individuals approach completion of their developmental requirements. Then, as days shorten, trees become increasingly responsive to cooling (Delpierre *et al*., 2009; Körner *et al*., 2016), strengthening the LST effect. Together, these changes can explain why, by August, trees responded similarly to cooling regardless of their earlier developmental trajectories (Fig. 1). These findings highlight the critical interaction between photoperiod and temperature in shaping the autumn phenology of *Fagus sylvatica*.

### Effects of daytime vs. nighttime temperature

In the second experiment we independently manipulated daytime and nighttime temperatures. The imposed decoupling of daytime and nighttime temperatures was not intended to represent common meteorological conditions. Rather, it was used to isolate processes that are inherently diel in nature. In trees, cell division and expansion predominantly occur at night (Steppe *et al*., 2015; Mencuccini *et al*., 2017; Zweifel et al., 2021), whereas carbon assimilation occurs primarily during the day (Fig. 6a). By experimentally separating these phases, we tested whether phenological responses depend on when within the diel cycle temperature constraints are applied. The strong and opposing responses observed under these forced asymmetries indicate that diel timing is a critical axis of physiological control, even if natural temperature cycles rarely produce such extreme contrasts.

**Figure 6.**
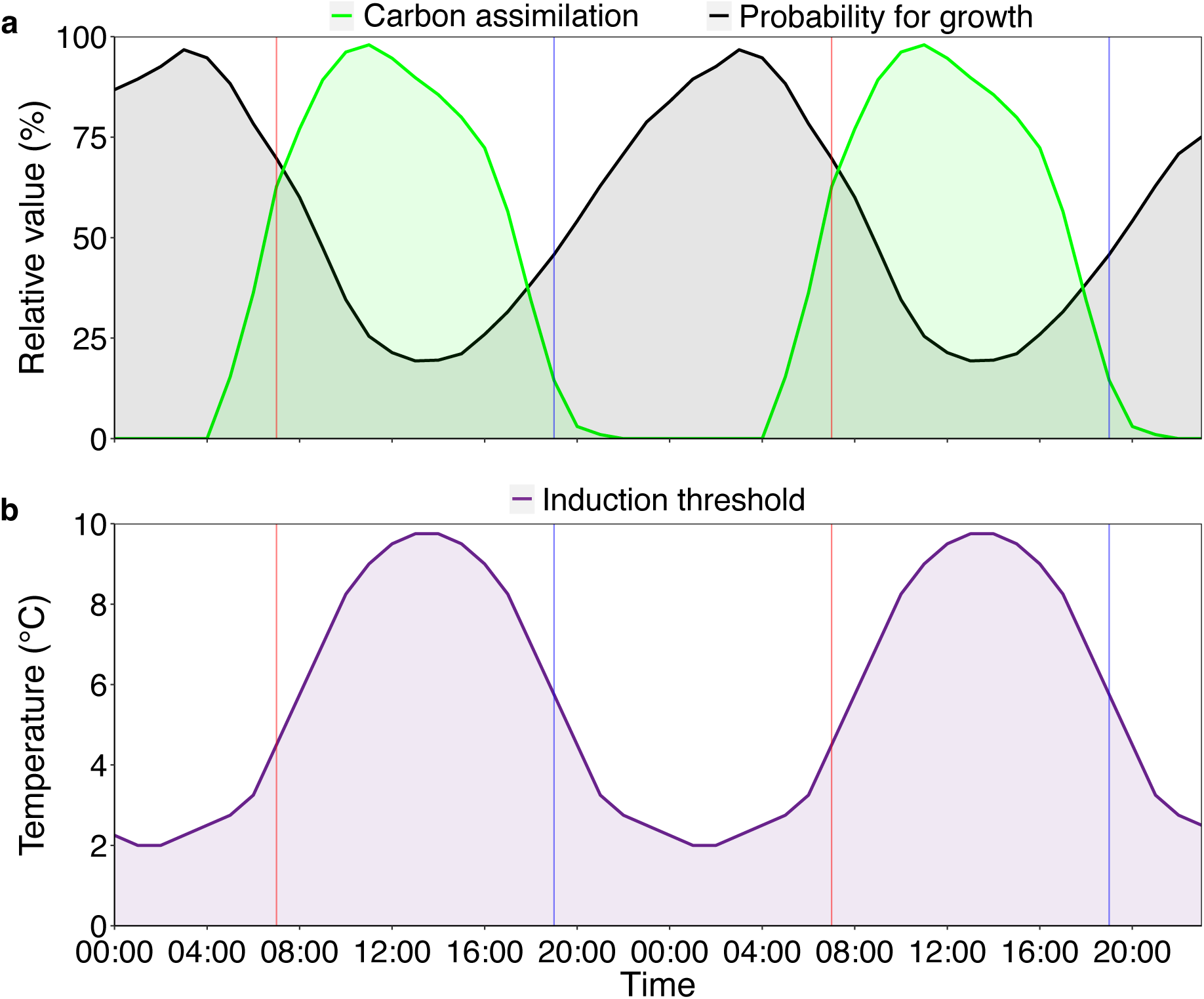
Diel patterns of relative growth, photosynthetic rate and theoretical thresholds for cold-induced bud set in *Fagus sylvatica*. Vertical red lines indicate the start of the day, and vertical blue lines indicate the start of the night, marking the boundaries between the 12-hour treatment windows used in experiment 2. **a)** The green curve shows relative carbon assimilation rate, the raw values were taken from the literature in a study that measured assimilation rates under controlled conditions (Urban *et al*., 2014). Values were linearly interpolated between measurements then converted to a percentage of the peak value. Finally, the curve was smoothed by taking the running mean of the target value, the previous value and the following value. The black curve shows the relative probability for growth, the raw values were taken from the literature in a study that measured growth rates in the field (Zweifel *et al*., 2021), then processed in the same way as the assimilation data. At night, low temperatures slow the trees’ developmental processes, possibly leading to delayed tissue maturation, while low temperatures during the day reduce photosynthetic activity. **b) The Diel Cooling Hypothesis:** As days shorten after the solstice, autumn bud set becomes increasingly responsive to cooling. Temperatures below a certain threshold induce overwintering responses, advancing autumn phenology. Our results indicate that daytime cooling of 8°C is below this threshold. However, because daily temperatures reach their minimum during the night, trees’ induction thresholds should be lower at night than during the day. Post-solstice night-time cooling of 8°C may thus have delayed bud set by slowing development rather than inducing overwintering responses.

Our experiments showed that daytime and nighttime temperatures had different effects on autumn phenology before and after the summer solstice: before the solstice, daytime cooling had no impact on the timing of bud set, while nighttime cooling and full-day cooling delayed bud set by 4.2 and 4.1 days, respectively (Fig. 5). This pattern was similar when using leaf senescence as the end-of-season marker. Because experimental cooling reduced photosynthesis by 69-83% across all treatments (Fig. S12), these effects are unlikely to be explained by changes in carbon assimilation. Instead, they highlight the importance of developmental processes—such as cell division and expansion—which mostly occur at night (Steppe *et al*., 2015; Mencuccini *et al*., 2017; Zweifel et al., 2021; Fig. 6a). The developmental arrest induced by night cooling—and therefore also by full-day cooling—likely slowed growth and maturation, effectively extending the growing season (ESD effect).

After the solstice, daytime and nighttime cooling of 8°C elicited opposite responses in both phenological markers, with weaker effects on senescence. Trees subjected to post-solstice daytime cooling and full-day cooling set their buds earliest, on average more than five days earlier than controls. As days shorten, the temperature threshold inducing this growth cessation must be lower at night, otherwise trees would risk prematurely ending their growing season due to the daily minimum temperatures occurring at night (Fig. 6b). This may explain why post-solstice nighttime cooling delayed, rather than advanced, autumn phenology: cooling of 8°C was sufficient to trigger overwintering responses—growth cessation and maturation of perennating tissues—when applied during the day but insufficient when applied at night (see Fig. S13 for a conceptual schematic).

An alternative explanation is that daytime cooling indirectly triggered dormancy induction by suppressing photosynthesis (Fig 6a), under the assumption that trees prioritise dormancy once photosynthesis is no longer possible (Körner *et al*., 2016). Though, as discussed above, late-season photosynthesis appears to assert little control over the autumn phenology of *F. sylvatica* (Zohner *et al*., 2023). Taken together, these results support a Diel Cooling Hypothesis, in which the impact of cooling depends on whether it occurs during the day or at night, because the temperature thresholds that trigger growth cessation differ between daytime and nighttime conditions.

Nighttime cooling always led to a delay in bud set of approximately four days (Fig. 5). As trees mainly grow at night (Steppe *et al*., 2015; Mencuccini *et al*., 2017; Zweifel *et al*., 2021), colder nighttime temperatures likely slowed down key developmental processes such as meristematic activity, tissue expansion and maturation, which in turn delayed primary growth cessation (Fig. 6). Similar responses may occur in other temperate tree species: cold autumn nights have been shown to delay growth cessation and slow bud development in *Populus, Pinus* and *Picea* species (Kramer, 1956; Malcolm & Pymar, 1975; Kalcsits *et al*., 2009). Moreover, the reversal in responses to temperature after the summer solstice has been observed consistently across Northern Hemisphere temperate forests (Zohner *et al*., 2023). Further experiments should explicitly test these diel responses in other temperate tree species as well as other provenances of beech.

### Support for the Solstice-as-Phenology Switch Hypothesis

The Solstice-as-Phenology-Switch hypothesis posits that temperature effects on autumn phenology reverse around the summer solstice, but does not assume that this reversal occurs exactly on June 21 (Zohner *et al*., 2023). Instead, the solstice represents the earliest point at which such a reversal can emerge, marking the onset of declining daylengths that enable phenological sensitivity to late-season cooling. Here we show how the antagonistic influences of the ESD effect and the LST effect can jointly determine the timing of this reversal.

Under this framework, the LST effect is ‘switched-on’ after the solstice as daylength begins to decline (Fig. 1). However, the observed reversal in temperature responses occurs only once early-season developmental effects (ESD effect) are balanced by increasing sensitivity to cooling (LST effect)—that is, at a compensatory point. Therefore, the timing of this reversal is not fixed to a calendar date but varies with developmental progression. Our conceptual model (Fig. 1) explicitly incorporates this flexibility, showing that the timing of the reversal depends on realised early-season development (ESD effect): under conditions that slow development (e.g., late leaf-out), the compensatory point is reached later in the season, whereas faster development advances it.

Our experiments support this framework: pre-solstice full-day cooling delayed bud set, whereas post-solstice full-day cooling advanced it, with differences between early– and late-developing individuals consistent with model predictions. Moreover, the contrasting impacts of daytime vs. nighttime cooling demonstrate that diel conditions further shape when the reversal is expressed. Thus, our findings support the Solstice-as-Phenology-Switch hypothesis and extend it by showing that its flexibility arises from interactions between developmental progression, diel temperature responses, and photoperiod.

Having established these effects under controlled physiological forcing, we next consider how the underlying mechanisms may operate under natural, fluctuating conditions. While beech trees are unlikely to experience sustained temperature regimes equivalent to the imposed chamber conditions, analogous developmental constraints can arise during cold springs, late cold spells following budburst, or at high-elevation and continental sites where temperatures remain low despite increasing photoperiod. Moreover, because developmental progression integrates temperature cumulatively over time, even short episodes of strong cooling can exert lasting carry-over effects on seasonal timing (Chuine, 2000; Liu *et al*., 2024). The relevance of our findings therefore lies not in the specific temperature regimes applied, but in identifying the processes through which temperature and development interact to shape autumn phenology.

Our results demonstrate that accelerating developmental progression—particularly via nighttime temperature effects—can advance the timing at which cooling begins to trigger growth cessation and senescence in European beech. This provides a mechanistic framework for interpreting phenological responses under climate warming, rather than making direct quantitative predictions under specific climate scenarios. These findings highlight the importance of accounting for developmental context and diel temperature sensitivity in phenological models of temperate trees.

## Acknowledgments

C.M.Z. was supported by the SNF Ambizione Fellowship programme (no. PZ00P3_193646) and T.W.C. by DOB Ecology and the Bernina Foundation.

## Author contributions

D.R. conceived and developed the study, conducted the experiments and analyses, and wrote the manuscript. C.M.Z equally contributed to the conception and development of the study, contributed to the experiments and analyses, and the text. L.M., H.M., Z.W. and Y.Z. contributed to the conception and implementation of the first experiment. R.B. contributed to the conception and implementation of the second experiment. T.W.C. and S.S.R. contributed to the text. All authors provided comments and approved the final manuscript.

## Data availability

The code and data for this study are available at Zenodo (Rebindaine *et al*., 2025).

## Supporting Information

**Figure S1.**
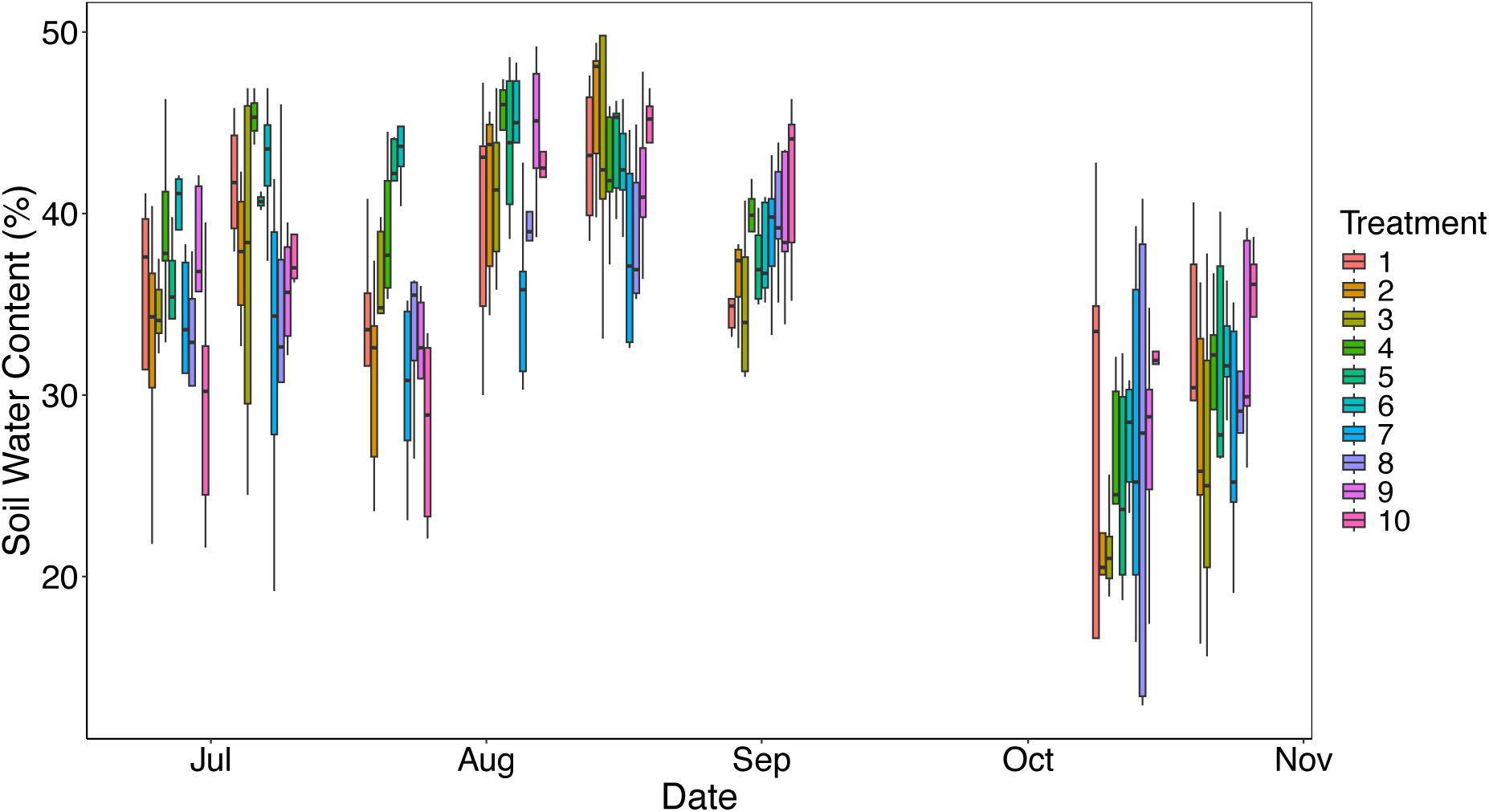
Seasonal soil water content (%) for each treatment (experiment 1). Five individuals per treatment were measured at each time step. All efforts were made to avoid water deficit and maintain equitable water availability across treatments. Across the experiment soil water content only significantly differed between treatments 6 and 7 (Tukey HSD test 7-6, estimate = –6%, p < 0.05). Differences in the responses between this treatment pair are not discussed in this study.

**Figure S2.**
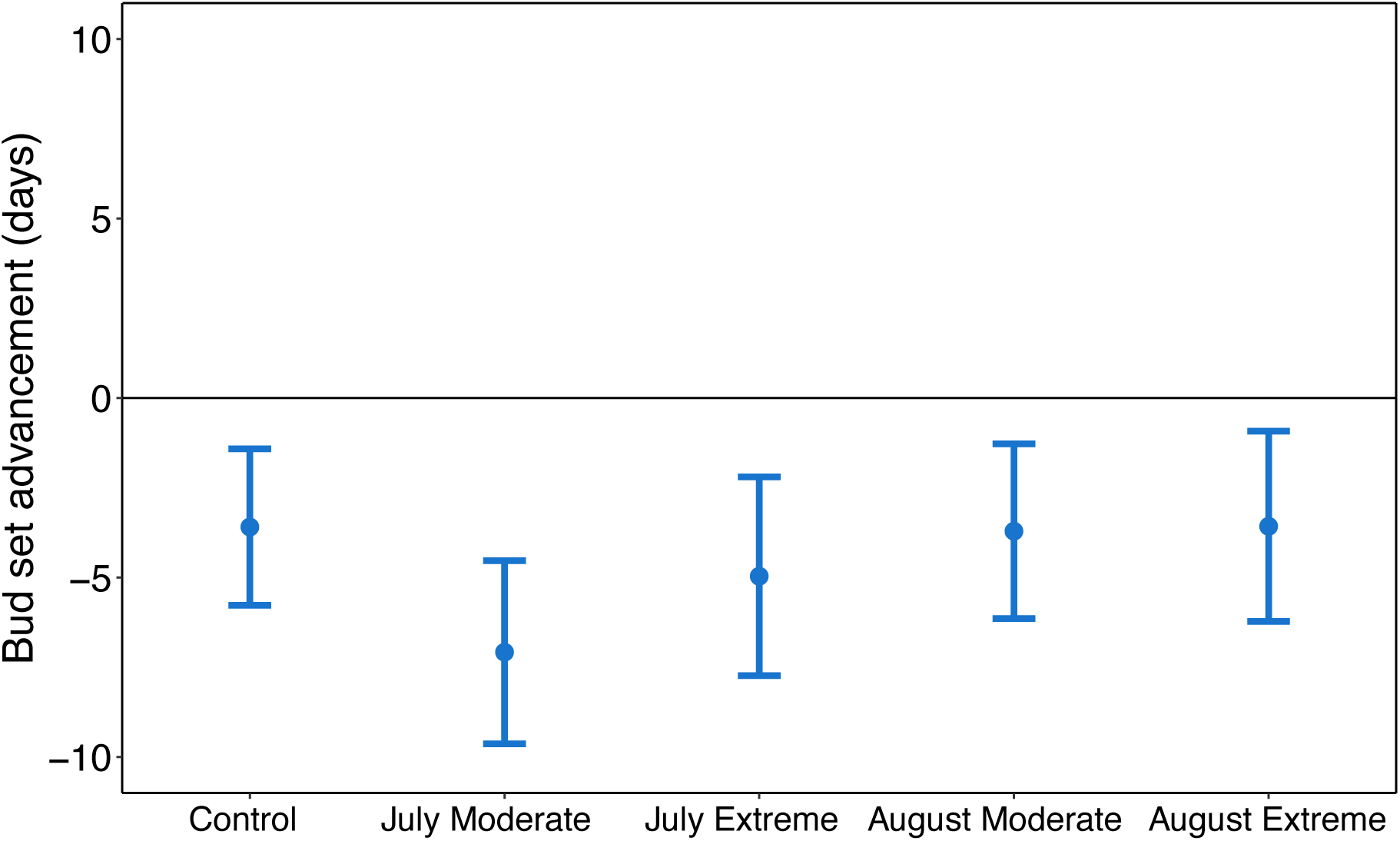
Effects of slow spring development on the timing of autumn bud set in *Fagus sylvatica* (experiment 1). Spring development was slowed by cooling the late-leafing trees from 4 April to 24 May in climate chambers. The late-leafing trees experienced a simulated day-night cycle of temperature and light availability, with minimum (2°C) and maximum (7°C) temperatures reached during the night– and daytime, respectively and otherwise simulated ambient conditions. Analyses show effect size means ± 95% confidence intervals from linear models, including treatment and bud-type (apical vs lateral) as predictors. Early-leafing effects are calculated against the early-leafing control and late-leafing effects are calculated against the late-leafing control. The bud type effect is not shown.

**Figure S3.**
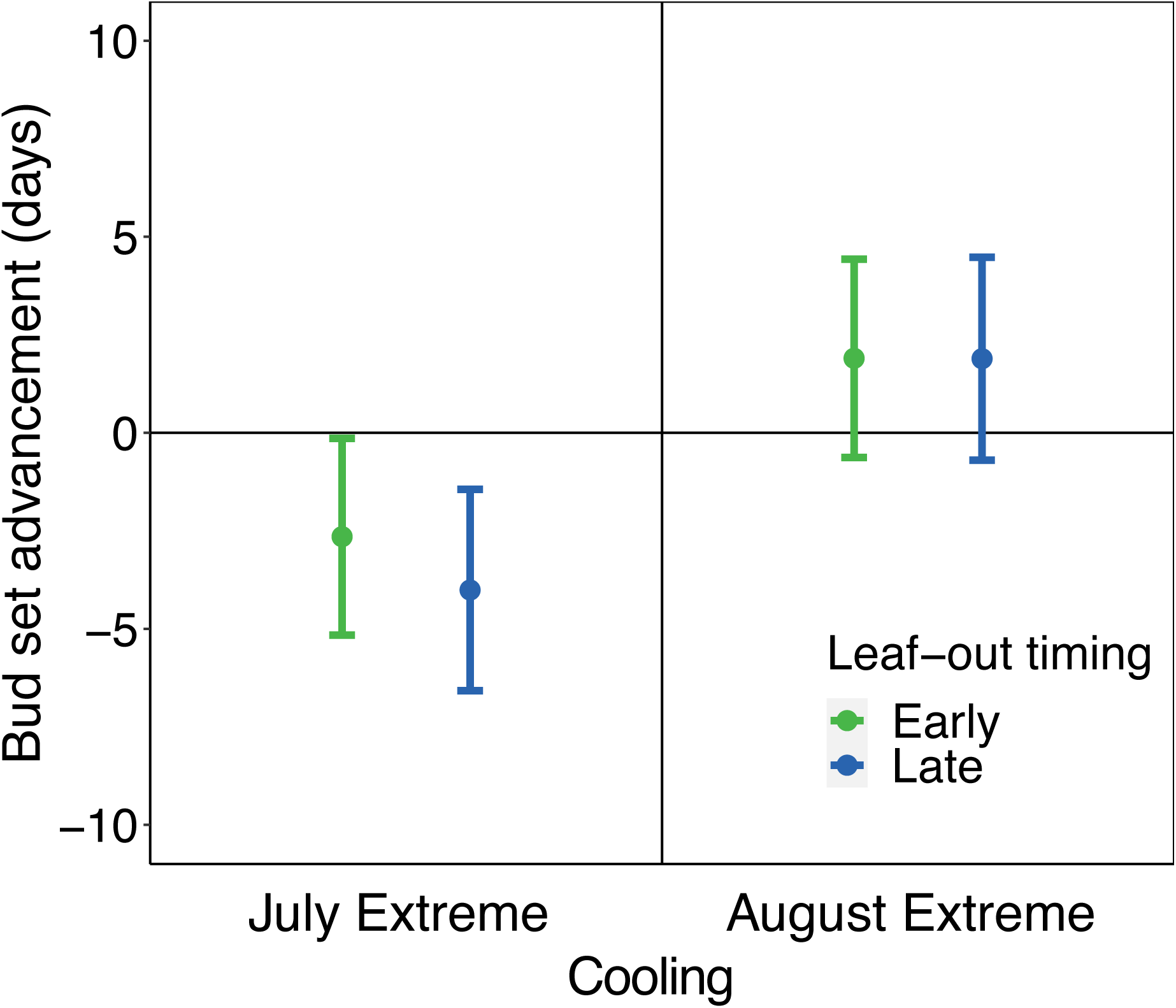
Effects of early-season development and extreme late-season cooling on the timing of autumn primary growth cessation in *Fagus sylvatica* (experiment 1). Effects of July (22 June to 23 July) and August (24 July to 25 August) extreme cooling (2-7°C) on bud set date for early– (green) and late– (blue) leafing trees. Analyses show effect size means ± 95% confidence intervals from linear models, including treatment and bud-type (apical vs lateral) as predictors. Early-leafing effects are calculated against the early-leafing control and late-leafing effects are calculated against the late-leafing control. The bud type effect is not shown.

**Figure S4.**
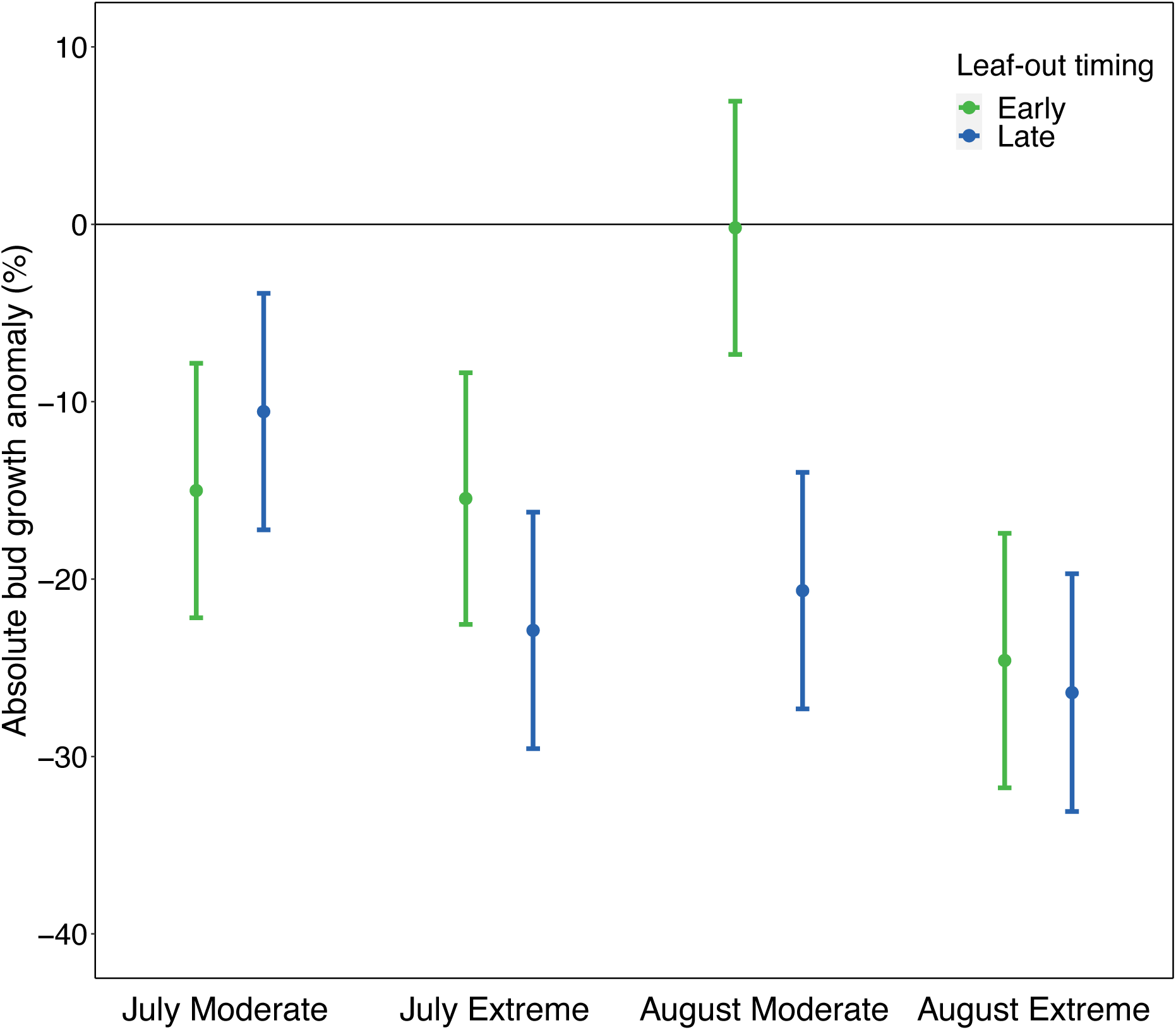
Effects of early-season development and late-season cooling on total bud growth in *Fagus sylvatica* (experiment 1). Effects of July (22 June to 23 July) and August (24 July to 25 August) moderate and extreme cooling (8-13°C and 2-7°C, respectively) on total seasonal (4 July to bud set date) bud growth for early– (green) and late– (blue) leafing trees. Analyses show effect size means ± 95% confidence intervals from linear models, including treatment and bud-type (apical vs lateral) as predictors. Early-leafing effects are calculated against the early-leafing control and late-leafing effects are calculated against the late-leafing control. The bud type effect is not shown.

**Figure S5.**
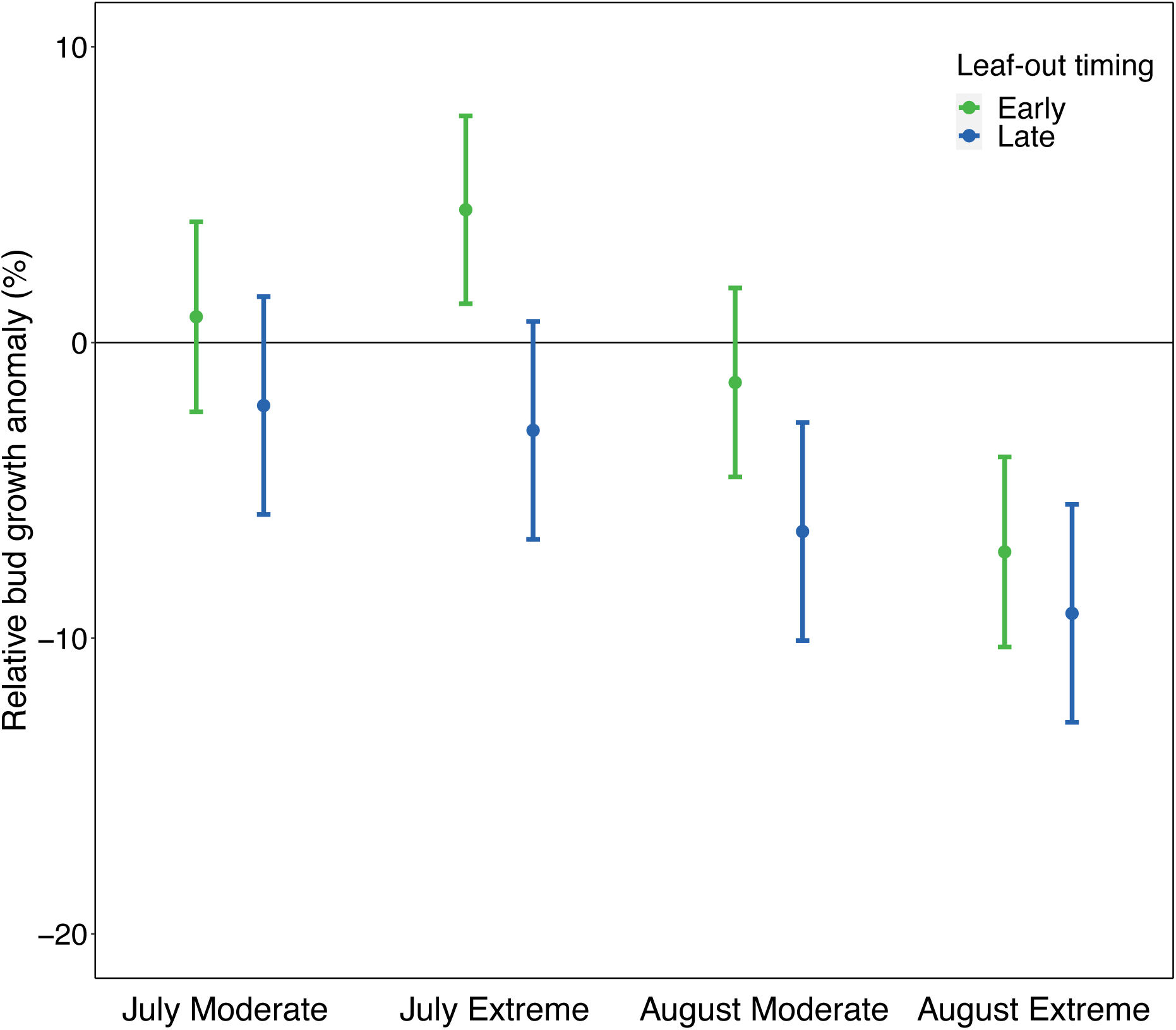
Effects of early-season development and late-season cooling on relative bud growth in *Fagus sylvatica* (experiment 1). Effects of July (22 June to 23 July) and August (24 July to 25 August) moderate and extreme cooling (8-13°C and 2-7°C, respectively) on relative seasonal (4 July to bud set date) bud growth for early– (green) and late– (blue) leafing trees. Analyses show effect size means ± 95% confidence intervals from linear models, including treatment and bud-type (apical vs lateral) as predictors. Early-leafing effects are calculated against the early-leafing control and late-leafing effects are calculated against the late-leafing control. The bud type effect is not shown.

**Figure S6.**
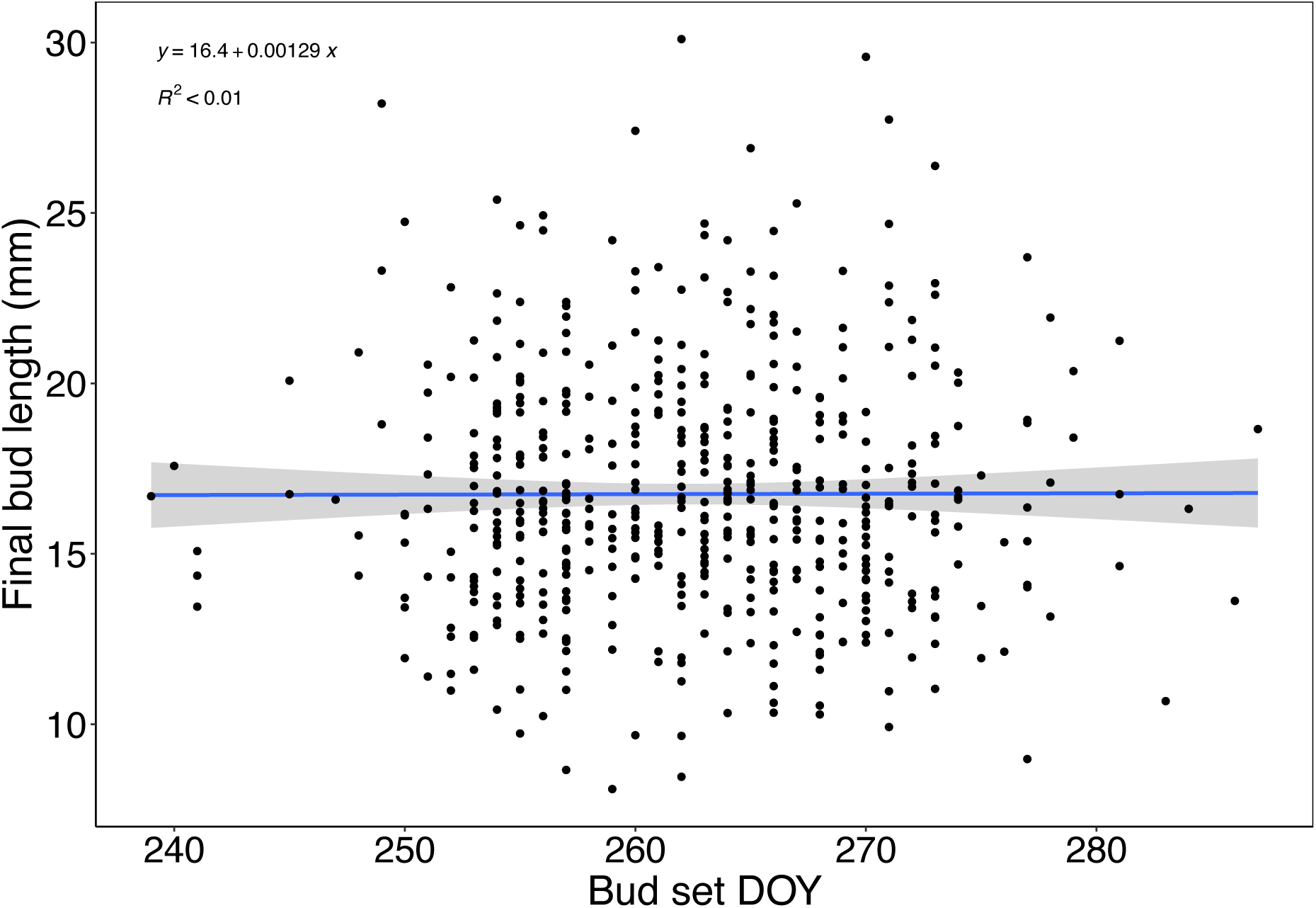
Relationship between end of season bud length and the timing of primary growth cessation in *Fagus sylvatica* (experiment 1). Effect of bud set DOY (day-of-year) on the final bud length at the end of the growing season from a linear model with 95% confidence interval.

**Table S1.** Mean and 95% confidence intervals of 50% leaf-out (day-of-year BBCH15), bud set (day-of-year), leaf-off (day-of-year 50% loss of leaf chlorophyll) and absolute bud growth (cm) for each treatment in experiment 1.

| TREATMENT | LEAF-OUT DOY<br>(95% CI) | BUD SET DOY<br>(95% CI) | LEAF-OFF DOY<br>(95% CI) | ABSOLUTE BUD<br>GROWTH CM (95%<br>CI) |
| --- | --- | --- | --- | --- |
| **1** | **131.6**<br>(130.2-133.0) | **260.5**<br>(259.1-261.9) | **317.3**<br>(314.5-320.1) | **12.88**<br>(11.96-13.80) |
| **2** | **148.1**<br>(147.8-148.4) | **264.1**<br>(262.4-265.8) | **322.4**<br>(320.2-324.6) | **14.21**<br>(13.53-14.89) |
| **3** | **131.2**<br>(129.7-132.7) | **261.9**<br>(260.0-263.8) | **319.6**<br>(317.8-321.4) | **10.95**<br>(10.29-11.61) |
| **4** | **148.6**<br>(148.2-149.0) | **268.9**<br>(267.2-270.6) | **325.5**<br>(323.5-327.5) | **12.72**<br>(11.99-13.45) |
| **5** | **130.1**<br>(128.9-131.3) | **263.1**<br>(261.1-265.1) | **322.8**<br>(320.5-325.1) | **10.89**<br>(10.41-11.37) |
| **6** | **148.7**<br>(148.4-149.0) | **268.1**<br>(266.2-270.0) | **323.4**<br>(321.7-325.1) | **10.97**<br>(10.35-11.59) |
| **7** | **130.2**<br>(128.6-131.8) | **256.0**<br>(254.6-257.4) | **312.2**<br>(309.4-315.0) | **12.85**<br>(12.17-13.53) |
| **8** | **148.9**<br>(148.6-149.2) | **259.7**<br>(257.7-261.7) | **319.8**<br>(318.3-321.3) | **11.28**<br>(10.63-11.93) |
| **9** | **132.3**<br>(130.8-133.8) | **258.6**<br>(256.5-260.7) | **319.1**<br>(316.6-321.6) | **9.71**<br>(9.22-10.20) |
| **10** | **148.0**<br>(147.6-148.4) | **262.2**<br>(260.5-263.9) | **320.0**<br>(317.9-322.1) | **10.46**<br>(9.81-11.11) |

**Table S2.** Mean and 95% confidence intervals of bud set (day-of-year), leaf-off (day-of-year 50% loss of leaf chlorophyll) and absolute bud growth (cm) for each treatment in experiment 2.

| TREATMENT | BUD SET DOY (95% CI) | LEAF-OFF DOY<br>(95% CI) | ABSOLUTE BUD<br>GROWTH CM (95% CI) |
| --- | --- | --- | --- |
| **AMBIENT**<br>**CONTROL** | **257.2** (254.7-259.7) | **321.1** (318.9-323.3) | **4.47** (3.80-5.14) |
| **PRE-**<br>**CHAMBER-**<br>**CONTROL** | **257.1** (254.1-260.1) | **319.8** (315.8-323.8) | **4.73** (3.82-5.54) |
| **PRE-FULL-**<br>**DAY** | **260.6** (257.3-263.9) | **322.7** (320.1-325.3) | **5.18** (4.26-6.10) |
| **PRE-DAY** | **257.2** (252.7-261.7) | **321.1** (318.4-323.8) | **4.95** (3.96-5.94) |
| **PRE-NIGHT** | **261.0** (257.9-264.1) | **322.8** (319.8-325.8) | **5.01** (4.34-5.68) |
| **POST-**<br>**CHAMBER-**<br>**CONTROL** | **255.4** (251.8-259.0) | **318.4** (315.8-321.0) | **4.24** (3.45-5.03) |
| **POST-FULL-**<br>**DAY** | **251.6** (248.3-254.9) | **320.9** (319.1-322.7) | **2.58** (2.08-3.08) |
| **POST-DAY** | **251.5** (248.0-255.0) | **316.3** (313.2-319.4) | **2.92** (2.32-3.52) |
| **POST-NIGHT** | **260.6** (256.7-264.5) | **320.6** (317.9-323.3) | **5.58** (4.70-6.46) |

**Figure S7.**
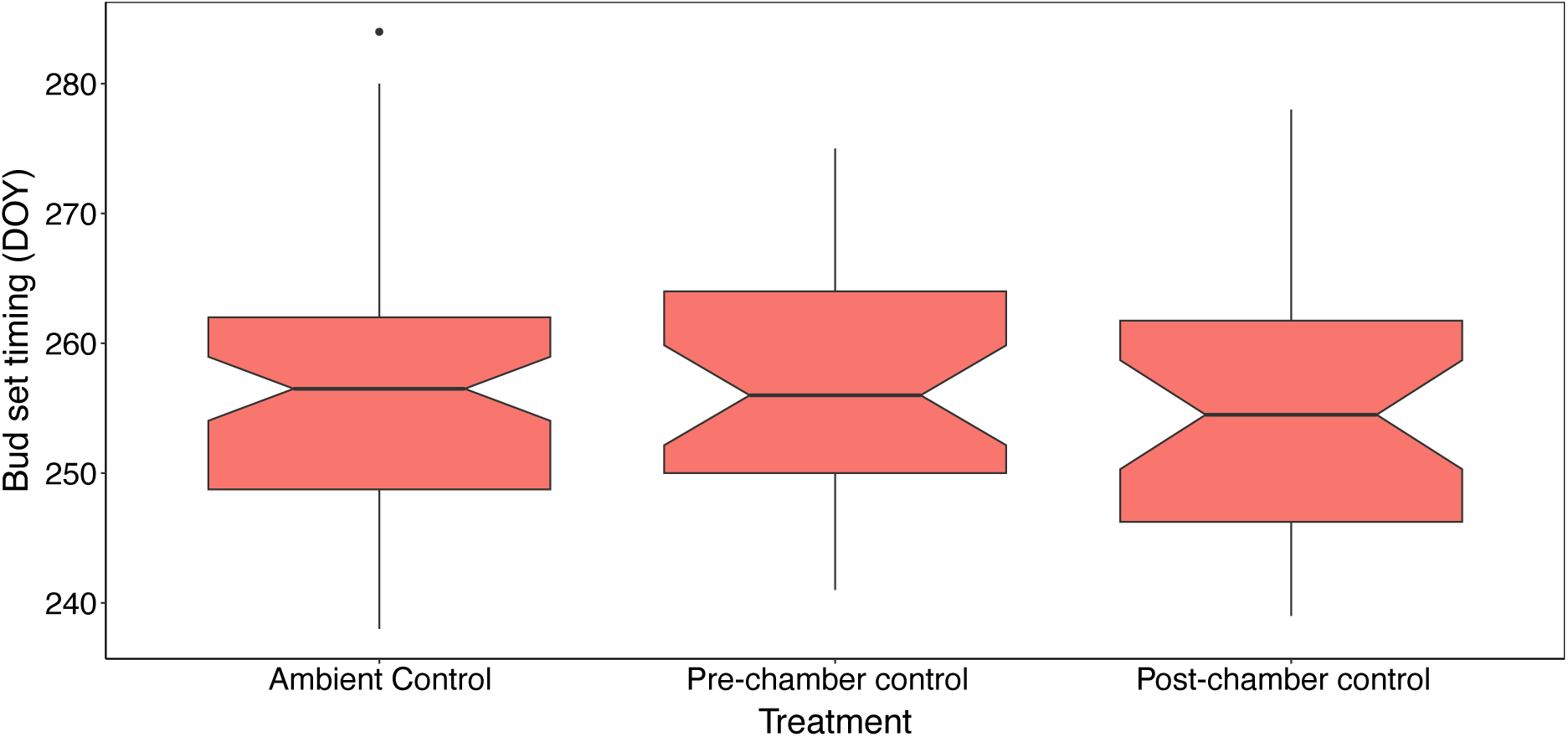
Bud set timing (day-of-year) for each control treatment in experiment 2. One-way ANOVA revealed no significant differences in bud set timing between the three control groups (*F*_2,136_ = 0.346, *p* = 0.708). Ambient control trees were placed outside throughout the experiment, Pre-Chamber control trees were in a climate chamber under simulated ambient conditions between 22 May and 21 June, Post-Chamber control trees were placed in a climate chamber under simulated ambient conditions between 22 June and 21 July.

**Figure S8.**
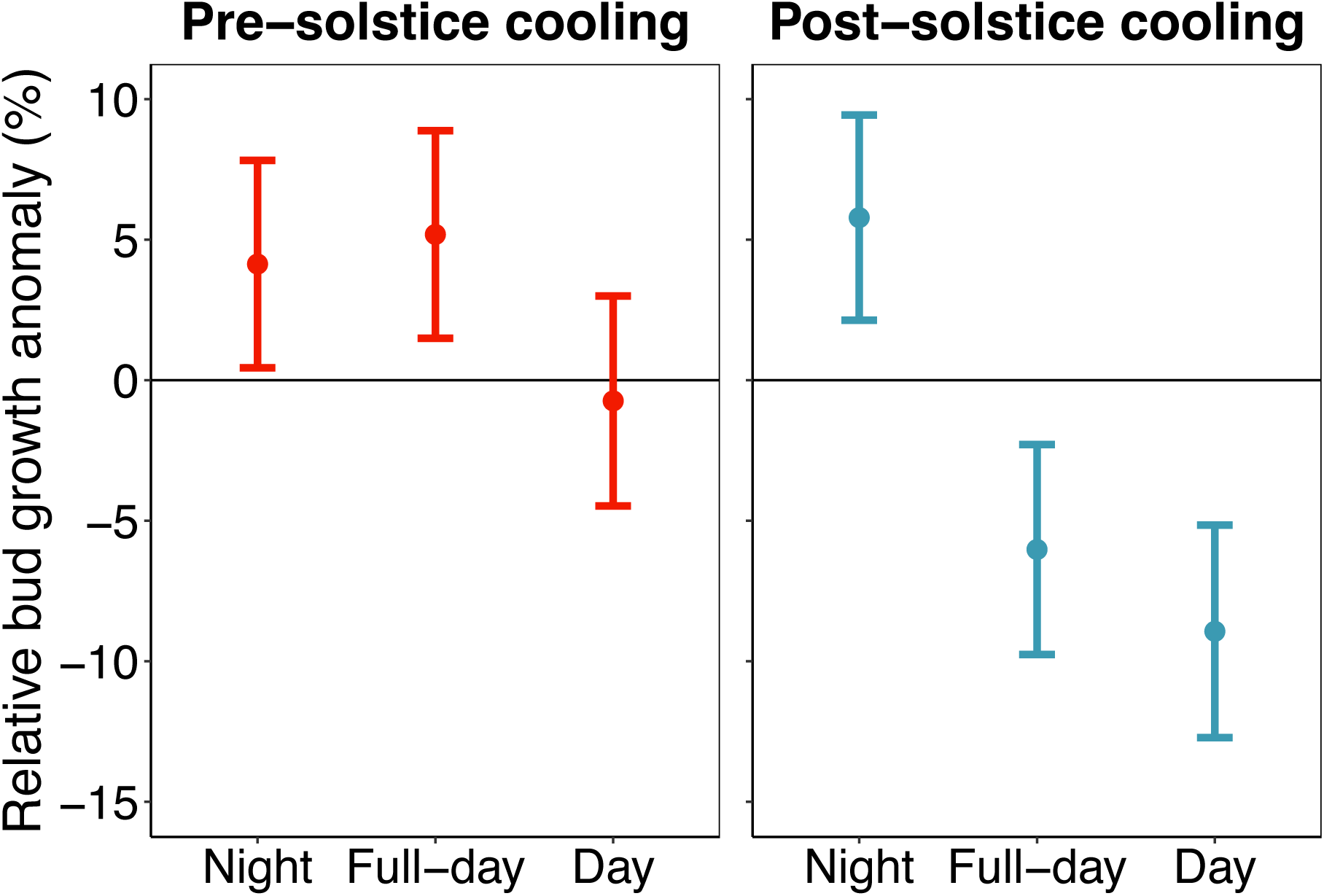
Effects of pre– and post-solstice daytime, nighttime and full-day cooling on relative bud growth in *Fagus sylvatica* (experiment 2). Pre-solstice (22 May to 21 June) and post-solstice (22 June to 21 July) cooling treatments were applied. Full-day cooling trees were continuously cooled to 8°C, day cooling trees were cooled to 8°C in the day and kept at 20°C at night, night cooling trees were cooled to 8°C at night and kept at 20°C in the day. Analyses show effect size means ± 95% confidence intervals from linear models, including treatment and bud-type (apical vs lateral) as predictors. The bud type effect is not shown.

**Figure S9.**
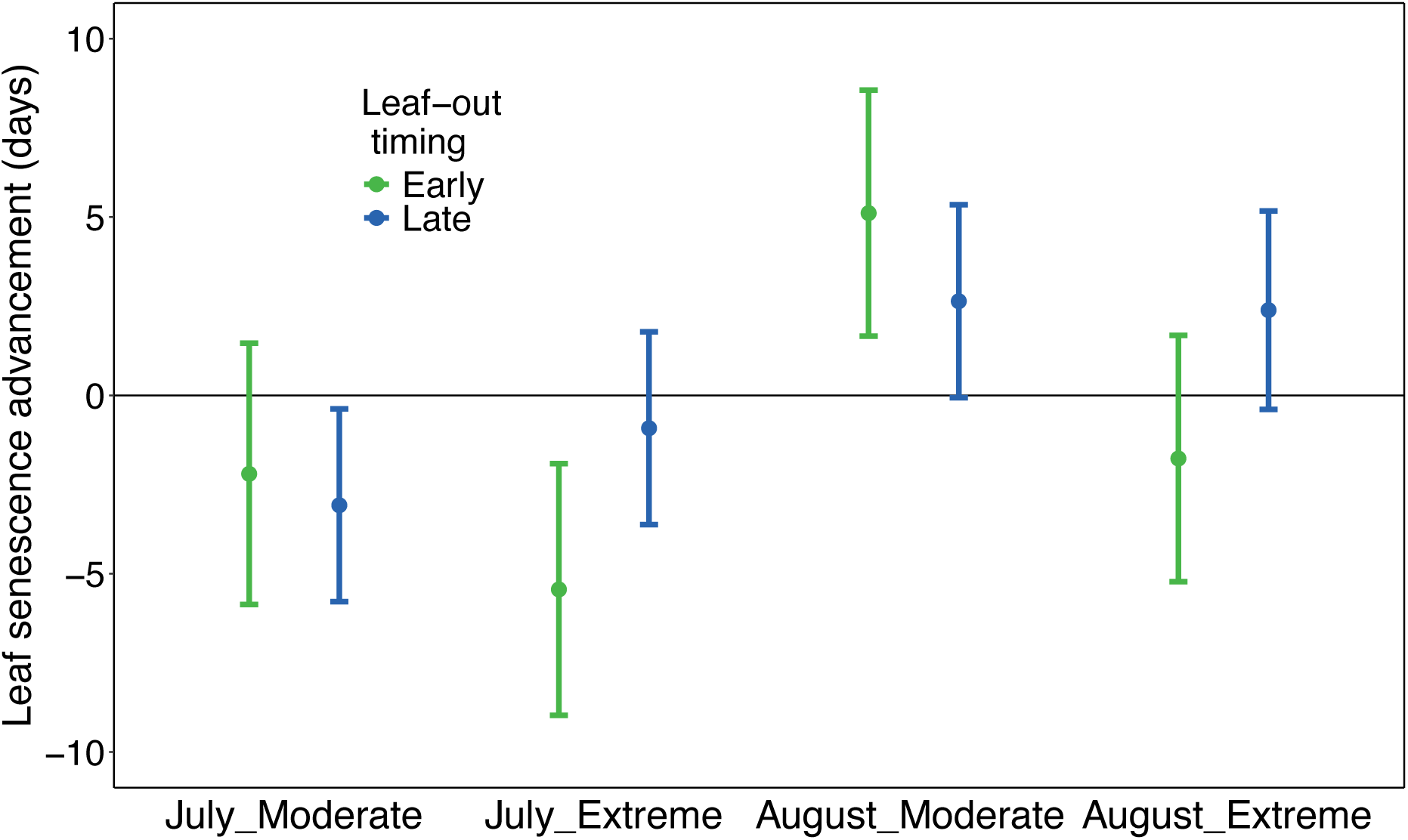
Effects of early-season development and late-season cooling on the timing of leaf senescence in *Fagus sylvatica* (experiment 1). Effects of July (22 June to 23 July) and August (24 July to 25 August) moderate (8-13°C) and extreme cooling (2-7°C) on date of 50% loss of leaf chlorophyll content for early– (green) and late– (blue) leafing trees. Analyses show effect size means ± 95% confidence intervals from linear models with treatment as the sole predictor. Early-leafing effects are calculated against the early-leafing control and late-leafing effects are calculated against the late-leafing control. Positive values indicate advances to leaf senescence and negative values indicate delays.

**Figure S10.**
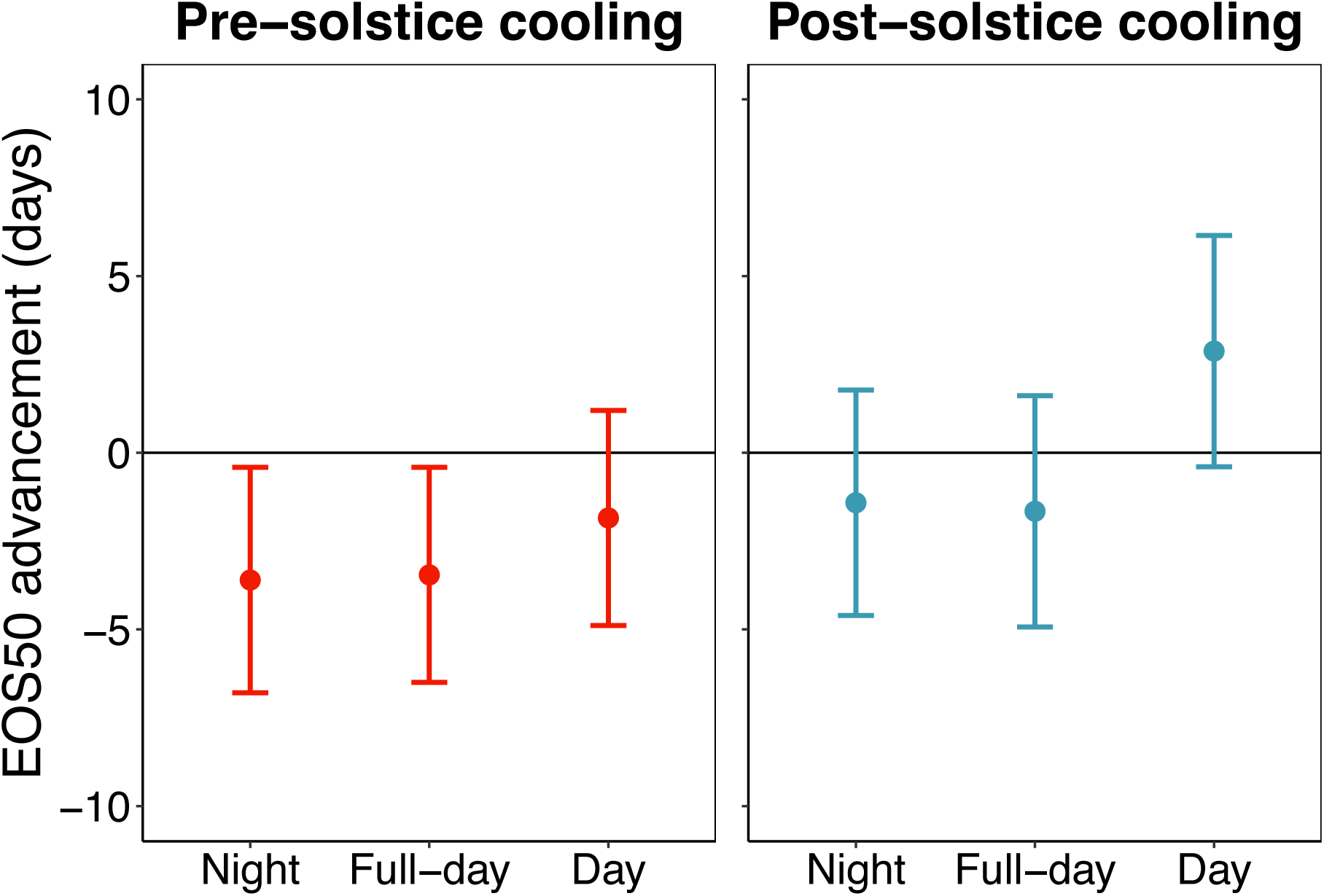
The pre– and post-solstice effects of night, full-day and day cooling on the timing of autumn leaf senescence in *Fagus sylvatica* (experiment 2). Effects of pre-solstice (22 May to 21 June) and post-solstice (22 June to 21 July) cooling on the date of 50% loss of leaf chlorophyll content. Full-day cooling trees were continuously cooled to 8°C, day cooling trees were cooled to 8°C in the day and kept at 20°C at night, night cooling trees were cooled to 8°C at night and kept at 20°C in the day. Analyses show effect size means ± 95% confidence intervals from linear models, including treatment as the sole predictor.

**Figure S11.**
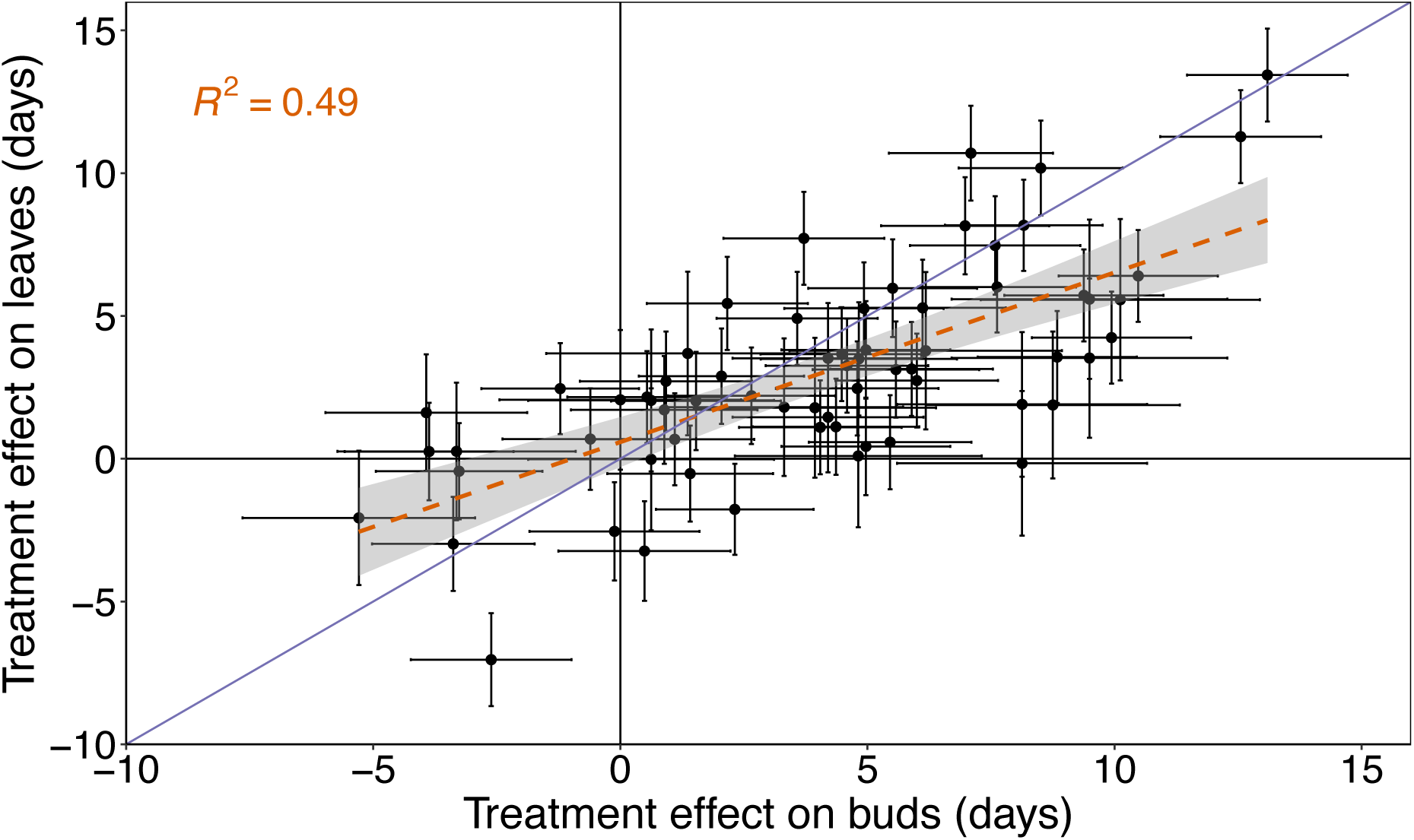
Treatment effects on the timing of bud set (x-axis) and leaf senescence (y-axis) across our two experiments. Each point represents the estimated treatment effect for a given contrast, with error bars showing standard errors. The dashed orange line indicates the fitted regression between the two metrics (*R^2^* = 0.49) along with the 95% confidence interval (shaded area), while the solid blue 1:1 line represents equal treatment effects on bud set and senescence. A majority of contrasts (83%) showed effects in the same direction for both phenometrics, indicating overall consistency in treatment impacts on the timing of autumn phenology.

**Figure S12.**
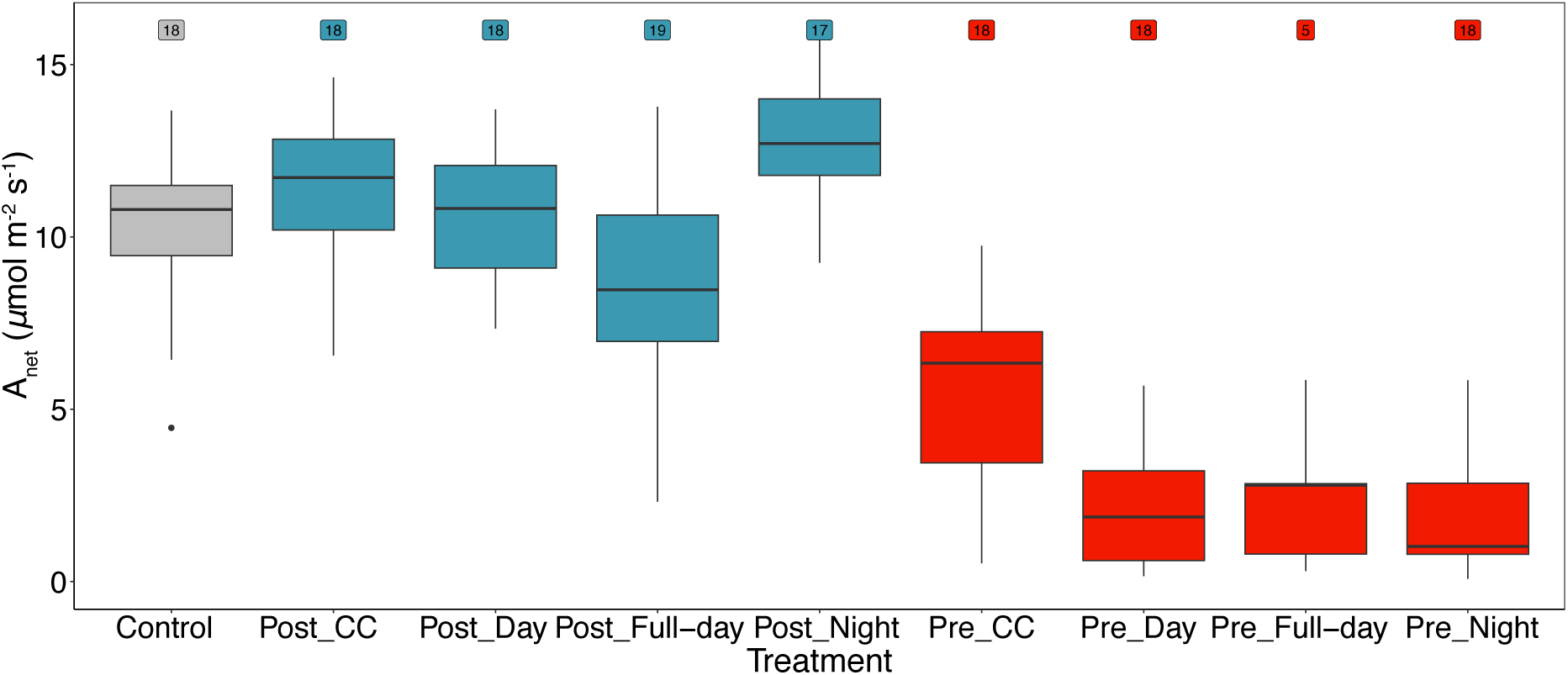
Photosynthesis rates of *Fagus sylvatica* trees during the pre-solstice treatment window (experiment 2). At this time, Control and post-solstice treatments (in blue) were outside under ambient conditions; pre-solstice treatments (in red) were in climate chambers under daytime, nighttime or full-day cooling or chamber control (CC) regimes. Absolute leaf-level photosynthesis (net CO_2_ assimilation). Insets at the top refer to the number of trees with valid assimilation data.

**Figure S13.**
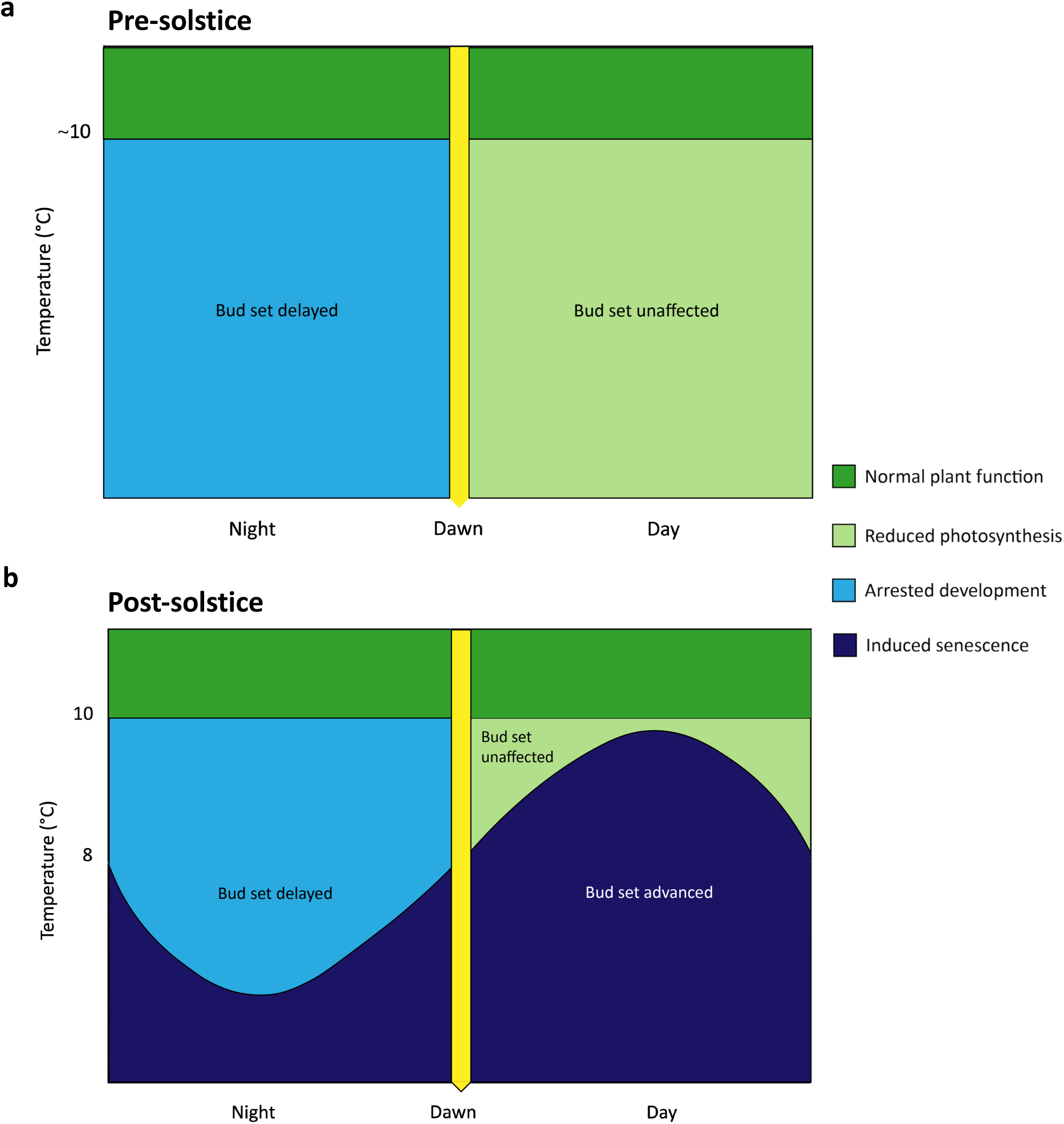
The Diel Cooling Hypothesis: Conceptual model of pre– and post-solstice effects of day and nighttime temperatures on the autumn phenology of *Fagus sylvatica*. The colours of the shaded areas represent the effect of cooling to different temperatures at specific times of the day. Above a certain threshold, plants function normally, and cooling has little effect on the timing of primary growth cessation (green; here depicted as 10°C, n.b. this is an intended simplification and does not necessarily accurately represent the reality). Light blue areas show combinations that lead to arrested development, which delays primary growth cessation. Pale green areas depict combinations that reduce photosynthesis and potentially growth but do not affect phenology. Dark purple areas depict combinations that lead to the induction of senescence and overwintering processes, therefore advancing primary growth cessation. **a)** Before the solstice, lengthening days give temperate trees a reliable cue of the favourable conditions to come, so there is no temperature that can induce senescence at this time of year (Kramer, 1936). Nighttime cooling (< ∼10°C) slows down beech’s developmental processes such as meristematic activity, tissue expansion and maturation, which leads to a growth and development debt (Zweifel *et al*., 2021). As the days are sufficiently warm, photosynthesis is unaffected. This means that the trees accumulate excess carbon that can only be used later in the growing season to make up for this developmental debt, thus delaying primary growth cessation. Daytime cooling (< ∼10°C) slows development to a lesser extent, however, it also reduces carbon gain by inhibiting photosynthesis but not respiration. Due to carbon limitation, affected trees may prioritise development over growth, so buds may be shorter but there is no resulting delay to primary growth cessation. **b)** After the solstice and due to shortening days, trees become increasingly sensitive to cooling, especially during the day. Therefore, daytime temperatures < ∼10°C are likely to induce overwintering responses and therefore advance autumn phenology. However, due to typical daily cycles of temperature always reaching their minimum during the night, trees are less susceptible to cooling in the night than in the day. At night, temperatures between ∼4°C and ∼10°C slow the trees’ developmental processes, leading to delayed primary growth cessation.

## References

1. Barbier FF, Dun EA, Beveridge CA. 2017. Apical dominance. Current Biology 27: R864–R865.

2. Bauerle WL, Oren R, Way DA, Qian SS, Stoy PC, Thornton PE, Bowden JD, Hoffman FM, Reynolds RF. 2012. Photoperiodic regulation of the seasonal pattern of photosynthetic capacity and the implications for carbon cycling. Proceedings of the National Academy of Sciences 109: 8612–8617.

3. Bezemer TM, Jones TH, Knight KJ. 1998. Long-term effects of elevated CO 2 and temperature on populations of the peach potato aphid Myzus persicae and its parasitoid Aphidius matricariae. Oecologia 116: 128–135.

4. Bigler C, Vitasse Y. 2021. Premature leaf discoloration of European deciduous trees is caused by drought and heat in late spring and cold spells in early fall. Agricultural and Forest Meteorology 307: 108492.

5. Boisvenue C, Running SW. 2006. Impacts of climate change on natural forest productivity – evidence since the middle of the 20th century: climate change impacts on forest vegetation. Global Change Biology 12: 862–882.

6. Buermann W, Forkel M, O’Sullivan M, Sitch S, Friedlingstein P, Haverd V, Jain AK, Kato E, Kautz M, Lienert S, et al. 2018. Widespread seasonal compensation effects of spring warming on northern plant productivity. Nature 562: 110–114.

7. Capdevielle-Vargas R, Estrella N, Menzel A. 2015. Multiple-year assessment of phenological plasticity within a beech (Fagus sylvatica L.) stand in southern Germany. Agricultural and Forest Meteorology 211-212: 13–22.

8. Carneros E, Yakovlev I, Viejo M, Olsen JE, Fossdal CG. 2017. The epigenetic memory of temperature during embryogenesis modifies the expression of bud burst-related genes in Norway spruce epitypes. Planta 246: 553–566.

9. Chuine I. 2000. A Unified Model for Budburst of Trees. Journal of Theoretical Biology 207: 337–347.

10. Cooke JEK, Eriksson ME, Junttila O. 2012. The dynamic nature of bud dormancy in trees: environmental control and molecular mechanisms. Plant, Cell & Environment 35: 1707–1728.

11. Crawley MJ, Akhteruzzaman M. 1988. Individual Variation in the Phenology of Oak Trees and Its Consequences for Herbivorous Insects. Functional Ecology 2: 409.

12. Cruz-García R, Balzano A, Čufar K, Scharnweber T, Smiljanić M, Wilmking M. 2019. Combining Dendrometer Series and Xylogenesis Imagery—DevX, a Simple Visualization Tool to Explore Plant Secondary Growth Phenology. Frontiers in Forests and Global Change 2: 60.

13. Delpierre N, Dufrêne E, Soudani K, Ulrich E, Cecchini S, Boé J, François C. 2009. Modelling interannual and spatial variability of leaf senescence for three deciduous tree species in France. Agricultural and Forest Meteorology 149: 938–948.

14. Dittmar C, Elling W. 2006. Phenological phases of common beech (Fagus sylvatica L.) and their dependence on region and altitude in Southern Germany. European Journal of Forest Research 125: 181–188.

15. Estiarte M, Peñuelas J. 2014. Alteration of the phenology of leaf senescence and fall in winter deciduous species by climate change: effects on nutrient proficiency. 21: 1005–1017.

16. Etzold S, Sterck F, Bose AK, Braun S, Buchmann N, Eugster W, Gessler A, Kahmen A, Peters RL, Vitasse Y, et al. 2022. Number of growth days and not length of the growth period determines radial stem growth of temperate trees (J Penuelas, Ed.). Ecology Letters 25: 427– 439.

17. Fu YSH, Campioli M, Vitasse Y, De Boeck HJ, Van den Berge J, AbdElgawad H, Asard H, Piao S, Deckmyn G, Janssens IA. 2014. Variation in leaf flushing date influences autumnal senescence and next year’s flushing date in two temperate tree species. Proceedings of the National Academy of Sciences 111: 7355–7360.

18. Fu YH, Piao S, Delpierre N, Hao F, Hänninen H, Liu Y, Sun W, Janssens IA, Campioli M. 2018. Larger temperature response of autumn leaf senescence than spring leaf-out phenology. Global Change Biology 24: 2159–2168.

19. Gessler A, Keitel C, Kreuzwieser J, Matyssek R, Seiler W, Rennenberg H. 2006. Potential risks for European beech (Fagus sylvatica L.) in a changing climate. Trees 21: 1–11.

20. Gessler A, Zweifel R. 2024. Beyond source and sink control – toward an integrated approach to understand the carbon balance in plants. New Phytologist 242: 858–869.

21. Gill AL, Gallinat AS, Sanders-DeMott R, Rigden AJ, Short Gianotti DJ, Mantooth JA, Templer PH. 2015. Changes in autumn senescence in northern hemisphere deciduous trees: a meta-analysis of autumn phenology studies. Annals of Botany 116: 875–888.

22. Kalcsits L, Kendall E, Silim S, Tanino K. 2009. Magnetic resonance microimaging indicates water diffusion correlates with dormancy induction in cultured hybrid poplar (Populus spp.) buds. Tree Physiology 29: 1269–1277.

23. Keenan TF, Gray JM, Friedl MA, Toomey M, Bohrer G, Hollinger DY, Munger JW, O’Keefe J, Schmid HP, Wing IS, et al. 2014. Net carbon uptake has increased through warming-induced changes in temperate forest phenology. Nature Climate Change.

24. Keenan TF, Richardson AD. 2015. The timing of autumn senescence is affected by the timing of spring phenology: implications for predictive models. Global Change Biology 21: 2634– 2641.

25. Körner C. 2006. Significance of Temperature in Plant Life. In: Morison JIL, Morecroft MD, eds. Plant Growth and Climate Change. Wiley, 48–69.

26. Körner C. 2021. Alpine Plant Life: Functional Plant Ecology of High Mountain Ecosystems. Cham: Springer International Publishing.

27. Körner C, Basler D, Hoch G, Kollas C, Lenz A, Randin CF, Vitasse Y, Zimmermann NE. 2016. Where, why and how? Explaining the low-temperature range limits of temperate tree species (M Turnbull, Ed.). Journal of Ecology 104: 1076–1088.

28. Körner C, Möhl P, Hiltbrunner E. 2023. Four ways to define the growing season. Ecology Letters 26: 1277–1292.

29. Kramer PJ. 1936. Effect of variation in length of day on growth and dormancy of trees. Plant Physiology 11: 127–137.

30. Kramer PJ. 1956. Some effects of various combinations of day and night temperatures and photoperiod on the height growth of loblolly pine seedlings. Forest Science 3: 45–55.

31. Krieger-Liszkay A, Krupinska K, Shimakawa G. 2019. The impact of photosynthesis on initiation of leaf senescence. Physiologia Plantarum 166: 148–164.

32. Lenz A, Vitasse Y, Hoch G, Körner C. 2014. Growth and carbon relations of temperate deciduous tree species at their upper elevation range limit (M Turnbull, Ed.). Journal of Ecology 102: 1537–1548.

33. Lim PO, Kim HJ, Gil Nam H. 2007. Leaf Senescence. Annual Review of Plant Biology 58: 115– 136.

34. Liu G, Migliavacca M, Reimers C, Kraft B, Reichstein M, Richardson AD, Wingate L, Delpierre N, Yang H, Winkler AJ. 2024. DeepPhenoMem V1.0: deep learning modelling of canopy greenness dynamics accounting for multi-variate meteorological memory effects on vegetation phenology. Geoscientific Model Development 17: 6683–6701.

35. Malcolm DC, Pymar CF. 1975. The influence of temperature on the cessation of height growth of Sitka spruce (Picea sitchensis Bong. Carr.). Silvae Genet 24: 5–6.

36. Malyshev AV, Van Der Maaten E, Garthen A, Maß D, Schwabe M, Kreyling J. 2022. Inter-Individual Budburst Variation in Fagus sylvatica Is Driven by Warming Rate. Frontiers in Plant Science 13: 853521.

37. Mariën B, Dox I, De Boeck HJ, Willems P, Leys S, Papadimitriou D, Campioli M. 2021. Does drought advance the onset of autumn leaf senescence in temperate deciduous forest trees? Biogeosciences 18: 3309–3330.

38. Mariën B, Robinson KM, Jurca M, Michelson IH, Takata N, Kozarewa I, Pin PA, Ingvarsson PK, Moritz T, Ibáñez C, et al. 2025. Nature’s Master of Ceremony: The Populus Circadian Clock as Orchestrator of Tree Growth and Phenology. npj Biological Timing and Sleep 2: 1–19.

39. Meger J, Ulaszewski B, Burczyk J. 2021. Genomic signatures of natural selection at phenology-related genes in a widely distributed tree species Fagus sylvatica L. BMC Genomics 22: 583.

40. Mencuccini M, Salmon Y, Mitchell P, Hölttä T, Choat B, Meir P, O’Grady A, Tissue D, Zweifel R, Sevanto S, et al. 2017. An empirical method that separates irreversible stem radial growth from bark water content changes in trees: theory and case studies. *Plant*, Cell & Environment 40: 290–303.

41. Menzel A, Sparks TH, Estrella N, Koch E, Aasa A, Ahas R, Alm-Kübler K, Bissolli P, Braslavská O, Briede A, et al. 2006. European phenological response to climate change matches the warming pattern: european phenological response to climate change. Global Change Biology 12: 1969–1976.

42. Nakagawa S, Westneat DF, Mizuno A, Araya-Ajoy YG, Class B, Dingemanse NJ, Dochtermann NA, Lagisz M, Laskowski KL, Pick JL, et al. 2026. Understanding different types of repeatability and intra-class correlation for an analysis of biological variation. Journal of the Royal Society Interface 23: 20250545.

43. Oksanen J, Simpson GL, Blanchet FG, Kindt R, Legendre P, Minchin PR, O’Hara RB, Solymos P, Stevens MHH, Szoecs E, et al. 2025. vegan: Community Ecology Package.

44. Paul MJ, Foyer CH. 2001. Sink regulation of photosynthesis. Journal of Experimental Botany 52: 1383–1400.

45. Peñuelas J, Rutishauser T, Filella I. 2009. Phenology Feedbacks on Climate Change. Science 324: 887–888.

46. Percival GC, Keary IP, Noviss K. 2008. The Potential of a Chlorophyll Content SPAD Meter to Quantify Nutrient Stress in Foliar Tissue of Sycamore (Acer pseudoplatanus), English Oak (Quercus robur), and European Beech (Fagus sylvatica). Arboriculture & Urban Forestry 34: 89–100.

47. Petterle A, Karlberg A, Bhalerao RP. 2013. Daylength mediated control of seasonal growth patterns in perennial trees. Current Opinion in Plant Biology 16: 301–306.

48. Piao S, Liu Q, Chen A, Janssens IA, Fu Y, Dai J, Liu L, Lian X, Shen M, Zhu X. 2019. Plant phenology and global climate change: Current progresses and challenges. Global Change Biology 25: 1922–1940.

49. Powell Graham R. 1988. Shoot elongation, leaf demography and bud formation in relation to branch position on Larix laricina saplings. Trees 2.

50. R Core Team. 2025. R: A Language and Environment for Statistical Computing.

51. Rebindaine D, Crowther TW, Susanne S. Renner, Zhaofei Wu, Yibiao Zou, Lidong Mo, Haozhi Ma, Raymo Bucher, Constantin M. Zohner. 2025. Developmental constraints mediate the summer solstice reversal of climate effects on the autumn phenology of European beech.

52. Richardson AD, Keenan TF, Migliavacca M, Ryu Y, Sonnentag O, Toomey M. 2013. Climate change, phenology, and phenological control of vegetation feedbacks to the climate system. Agricultural and Forest Meteorology 169: 156–173.

53. Rohde A, Bhalerao RP. 2007. Plant dormancy in the perennial context. Trends in Plant Science 12: 217–223.

54. Scotti I, González-Martínez SC, Budde KB, Lalagüe H. 2016. Fifty years of genetic studies: what to make of the large amounts of variation found within populations? Annals of Forest Science 73: 69–75.

55. Signarbieux C, Toledano E, Sanginés de Carcer P, Fu YH, Schlaepfer R, Buttler A, Vitasse Y. 2017. Asymmetric effects of cooler and warmer winters on beech phenology last beyond spring. Global Change Biology 23: 4569–4580.

56. Singh RK, Bhalerao RP, Eriksson ME. 2021. Growing in time: exploring the molecular mechanisms of tree growth. Tree Physiology 41: 657–678.

57. Singh RK, Bhalerao RP, Maurya JP. 2022. When to branch: seasonal control of shoot architecture in trees. The FEBS Journal 289: 8062–8070.

58. Singh RK, Svystun T, AlDahmash B, Jönsson AM, Bhalerao RP. 2017. Photoperiod– and temperature-mediated control of phenology in trees – a molecular perspective. New Phytologist 213: 511–524.

59. Solvin TM, Steffenrem A. 2019. Modelling the epigenetic response of increased temperature during reproduction on Norway spruce phenology. Scandinavian Journal of Forest Research 34: 83–93.

60. Steppe K, Sterck F, Deslauriers A. 2015. Diel growth dynamics in tree stems: linking anatomy and ecophysiology. Trends in Plant Science 20: 335–343.

61. Švik M, Brovkina O, Veljanovski T, Čater M. 2025. Phenological trends of European beech stands along the Carpathian arc: a 20-year MOD13Q1/MYD13Q1 based analysis. European Journal of Remote Sensing 58: 2506576.

62. Tanino KK, Kalcsits L, Silim S, Kendall E, Gray GR. 2010. Temperature-driven plasticity in growth cessation and dormancy development in deciduous woody plants: a working hypothesis suggesting how molecular and cellular function is affected by temperature during dormancy induction. Plant Molecular Biology 73: 49–65.

63. Thackeray SJ, Henrys PA, Hemming D, Bell JR, Botham MS, Burthe S, Helaouet P, Johns DG, Jones ID, Leech DI, et al. 2016. Phenological sensitivity to climate across taxa and trophic levels. Nature 535: 241–245.

64. Tumajer J, Kašpar J, Kuželová H, Shishov VV, Tychkov II, Popkova MI, Vaganov EA, Treml V. 2021. Forward Modeling Reveals Multidecadal Trends in Cambial Kinetics and Phenology at Treeline. Frontiers in Plant Science 12: 613643.

65. Urban O, Klem K, Holišová P, Šigut L, Šprtová M, Teslová-Navrátilová P, Zitová M, Špunda V, Marek MV, Grace J. 2014. Impact of elevated CO2 concentration on dynamics of leaf photosynthesis in Fagus sylvatica is modulated by sky conditions. Environmental Pollution 185: 271–280.

66. Vitasse Y, François C, Delpierre N, Dufrêne E, Kremer A, Chuine I, Delzon S. 2011. Assessing the effects of climate change on the phenology of European temperate trees. Agricultural and Forest Meteorology 151: 969–980.

67. Vose RS, Easterling DR, Gleason B. 2005. Maximum and minimum temperature trends for the globe: An update through 2004. Geophysical Research Letters 32: 2005GL024379.

68. Yang LH, Rudolf VHW. 2010. Phenology, ontogeny and the effects of climate change on the timing of species interactions. Ecology Letters 13: 1–10.

69. Zani D, Zani D, Crowther TW, Mo L, Renner SS, Zohner CM. 2020. Increased growing-season productivity drives earlier autumn leaf senescence in temperate trees. Science.

70. Zhong Z, He B, Chen HW, Chen D, Zhou T, Dong W, Xiao C, Xie S, Song X, Guo L, et al. 2023. Reversed asymmetric warming of sub-diurnal temperature over land during recent decades. Nature Communications 14: 7189.

71. Zohner CM, Mirzagholi L, Renner SS, Mo L, Rebindaine D, Bucher R, Palouš D, Vitasse Y, Fu YH, Stocker BD, et al. 2023. Effect of climate warming on the timing of autumn leaf senescence reverses after the summer solstice. Science 381: eadf5098.

72. Zohner CM, Renner SS. 2019. Ongoing seasonally uneven climate warming leads to earlier autumn growth cessation in deciduous trees. Oecologia 189: 549–561.

73. Zweifel R, Sterck F, Braun S, Buchmann N, Eugster W, Gessler A, Häni M, Peters RL, Walthert L, Wilhelm M, et al. 2021. Why trees grow at night. New Phytologist 231: 2174–2185.

